# Cofilin Suppresses Tau-Induced Defects in Dense-Core Granule Formation and Aβ-Induced Neurodegeneration

**DOI:** 10.64898/2026.09.22.753479

**Authors:** Bhavna Verma, Amy Cording, Lewis Blincowe, Shauna Rice, Harrison Vine, S. Mark Wainwright, Deborah C I Goberdhan, Clive Wilson

## Abstract

Intracellular neurofibrillary tangles formed from hyperphosphorylated tau and extracellular amyloid plaques containing aggregated Aβ-peptides, specific cleavage products of the Amyloid Precursor Protein (APP), are the primary histopathological hallmarks of Alzheimer’s Disease (AD), the leading cause of dementia in humans. However, the initiating steps that lead to these pathologies and early neurodegeneration, and the mechanisms by which tau- and Aβ-induced effects might be linked remain unclear. Using the prostate-like secondary cell (SC) in *Drosophila*, we recently showed that Aβ modulates normal APP- and membrane-associated protein aggregation in the dense-core granule (DCG) compartments of the regulated secretory pathway by interfering with subsequent membrane:DCG dissociation. This disrupts endolysosomal trafficking and propagates the resulting endolysosomal defects to other cells that endocytose the secreted abnormal DCG proteins. Here we show that overexpressing human tau also disrupts DCG aggregation and membrane:DCG dissociation inside SC secretory compartments, leading to increased endolysosomal targeting of these compartments. In a genetic screen, we find that knockdown of *cofilin*, which encodes an actin-severing protein required for dynamic remodelling of microfilaments, generates a similar phenotype. Consistent with this, overexpression of Cofilin, which is known to suppress tau-induced neurodegeneration in flies, reduces tau-induced DCG defects in SCs. Indeed, we find that Cofilin overexpression also suppresses Aβ-induced degeneration in the fly eye. We conclude that membrane:DCG aggregate dissociation in DCG compartments is disrupted by both tau- and Aβ-induced genetic changes that are relevant to AD, and this partially involves inhibition of actin cytoskeleton dynamics. Increasing actin remodelling activity can suppress neurodegeneration induced by both tau and Aβ, suggesting that this process provides an important functional link between them that might be targeted therapeutically.

## Introduction

Alzheimer’s Disease (AD) is the leading cause of dementia worldwide and as lifespan continues to increase, its prevalence is predicted to rise, posing serious health and social care challenges (Gustavsson et al., 2023). At a cellular level, AD is characterised by two major hallmarks: first, extracellular amyloid plaques primarily composed of amyloid-β (Aβ) peptides, cleavage products of the Amyloid Precursor Protein (APP) (Weglinski and Jeans, 2023), and second, cytosolic neurofibrillary tangles containing hyperphosphorylated forms of microtubule-regulating cytoskeletal protein tau, encoded by the *MAPT* gene (Rawat et al., 2022).

Excess levels of Aβ and amyloid plaques can often precede onset of dementia by many years, which led to the proposal of the ‘amyloid hypothesis’ as an explanation of AD onset (Selkoe and Hardy, 2016). Ultimately, however, plaque burden is a less accurate predictor of disease severity than neurofibrillary tangles (Morris et al., 2014). Importantly, the coincidence of these two hallmarks strongly suggests that their emergence is connected, most likely via the intracellular events that initially generate Aβ.

Intracellular Aβ production involves APP cleavage by β-secretase, thought to occur primarily within recycling and early endosomal compartments (Buggia-Prévot et al., 2014; Das et al., 2016; Wang et al., 2024), followed by γ-secretase cleavage, which is most prevalent inside late endosomes and lysosomes (LELs) (Ni et al., 2006; Sannerud et al., 2016). Interestingly, abnormal endolysosomal trafficking is one of the earliest neuronal pathologies observed in patients, both with familial and more common sporadic forms of AD (Cataldo et al., 1996; Hung et al., 2018; Mishra et al., 2024; Krogsaeter et al., 2025). This has led to a revision of the amyloid hypothesis, the endolysosomal ‘traffic jam’ model (Kimura and Yanagisawa, 2018). In this model, accumulating intracellular Aβ and Aβ-oligomers, which are intermediates in the aggregation process that generates plaques, induce endolysosomal defects, particularly at the synapse, that trigger neurodegeneration.

Importantly, Aβ is also reported to drive tau pathology via a number of mechanisms involving endolysosomes. For example, in rodent models, accumulation of endocytosed Aβ inside neurons can induce tau hyperphosphorylation (Gao et al., 2025), while endolysosomal Aβ-oligomers promote pathological changes in the subcellular organisation of tau (Schützmann et al., 2021). Amyloid plaques appear to induce seeding of tau aggregation and intercellular propagation of these seeds in mouse models (He et al., 2018). Indeed, in human brain preparations, Aβ can promote transfer of pathological seeds of aggregated tau between cells in exosomes, extracellular vesicles (EVs) generated in endosomal compartments (Miyoshi et al., 2021; Fowler et al., 2025). These findings suggest interactions between Aβ and tau that disrupt the cytoskeleton and intracellular trafficking, promote cell-to-cell propagation of pathology, and might exacerbate Aβ-induced endolysosomal defects. However, because of the small size of endosomal compartments, it has been challenging to visualise the dynamic events driving such changes and how they might be triggered.

We have recently characterised a prostate-like secretory cell in *Drosophila melanogaster*, the secondary cell (SC), that has highly enlarged secretory and lysosomal compartments (Corrigan et al., 2014; Redhai et al., 2016; Fan et al., 2020). The maturation of secretory compartments can be imaged in these cells in real-time. In this system, regulated secretory compartments undergo a switch to recycling endosomal identity, marked by the small GTPase, Rab11; this is required for normal intra-compartmental aggregation of secreted proteins into insoluble, spherical dense-core granules (DCGs) and biogenesis of intraluminal vesicles (ILVs) that will be secreted as Rab11-exosomes (Fan et al., 2020; Wells et al., 2023).

These fundamental processes appear to be conserved in human cells (Stockhammer et al., 2024; Wang et al., 2025). Indeed, in mammalian, as well as fly, neurons, Rab11 isoforms also play a key role in regulated synaptic secretion (Khvotchev et al., 2003; Beronja et al., 2005).

Additional studies have revealed that *Drosophila* APP is required for normal DCG formation in SCs, a function that can be replaced by human APP (Singh et al., 2025). APP promotes membrane-primed protein aggregation and must then be cleaved, probably by β-secretase, for a normal large central DCG to form within Rab11-positive secretory compartments. Occasionally, these compartments appear to fail quality control and are acidified through interactions with small LELs, then fully degraded by fusion with the lysosome (Singh et al., 2025).

Expressing Aβ in SC secretory compartments disrupts membrane:aggregate dissociation during early maturation, increasing lysosomal targeting, but the defective secretory compartments lack normal motility and frequently fail to fuse with the lysosome (Singh et al., 2025). Some of these compartments appear to release their contents by fusing with the plasma membrane. The secreted DCG proteins are abnormally endocytosed by other cells, leading to their accumulation in defective enlarged LELs, thus propagating the endolysosomal defect (Singh et al., 2025, Cording et al., 2026). Therefore, in this cell type, early defects in DCG compartment maturation can result in endolysosomal trafficking defects that are exacerbated by increased Aβ formation and then propagated to other cells. A range of studies characterising the links between APP, Aβ, recycling endosomes and endolysosomes in neurons are consistent with a model in which these changes might trigger neurodegeneration (Verma et al., 2026).

Since tau may interplay with Aβ in generating cellular AD pathologies, we reasoned that it might also affect early events in DCG biogenesis. Here, we test this hypothesis in SCs and find that overexpressing human tau alters the earliest stages of DCG and ILV biogenesis, frequently generating a membrane-associated cylindrical DCG, and driving increased compartmental targeting to lysosomes. By studying the effects of knocking down other cytoskeletal regulators, we show that tau and actin-severing protein Cofilin, previously reported to suppress tau-induced neurodegeneration phenotypes in the eye (Fulga et al., 2007), antagonistically regulate normal DCG formation. Furthermore, increasing Cofilin levels suppresses the effects of Aβ in a *Drosophila* eye neurodegeneration model. We conclude that reducing actin remodelling activity plays a critical role downstream of both Aβ and tau in generating AD-relevant pathologies, potentially via a mechanism that disrupts regulated secretion and associated endolysosomal trafficking.

## Results

### Human tau-2N4R overexpression in SCs disrupts DCG protein aggregation and endolysosomal trafficking of the resulting defective DCG compartments

Overexpression of the longest form of human tau, htau-2N4R, in *Drosophila* neurons has previously been reported to induce neurotoxicity using a range of assays that detect, for example, photoreceptor degeneration in the eye, climbing defects and lifespan reduction (Wittmann et al., 2001; Jackson et al., 2002; Gorsky et al., 2016; Hannan et al., 2016).

We overexpressed htau-2N4R (hereon referred to as htau) in adult SCs under UAS-GAL4 control (Brand and Perrimon, 1993), using GAL80^ts^-regulated *dsx-GAL4*. This driver is exclusively expressed in the ∼40 SCs at the distal tip of each lobe within the bilobed male accessory gland (MAG; Fig. 1A) when flies are switched to 29°C at eclosion (Fan et al., 2020). The flies also harboured a GFP-tagged version of MFAS, the orthologue of human extracellular matrix protein, Transforming Growth Factor-Induced (TGFBI), expressed from the endogenous gene locus; each SC contains approximately ten DCG compartments and this protein normally concentrates in DCG aggregates within them (Singh et al., 2025; Fig. 1A and 1B).

**Fig. 1.**
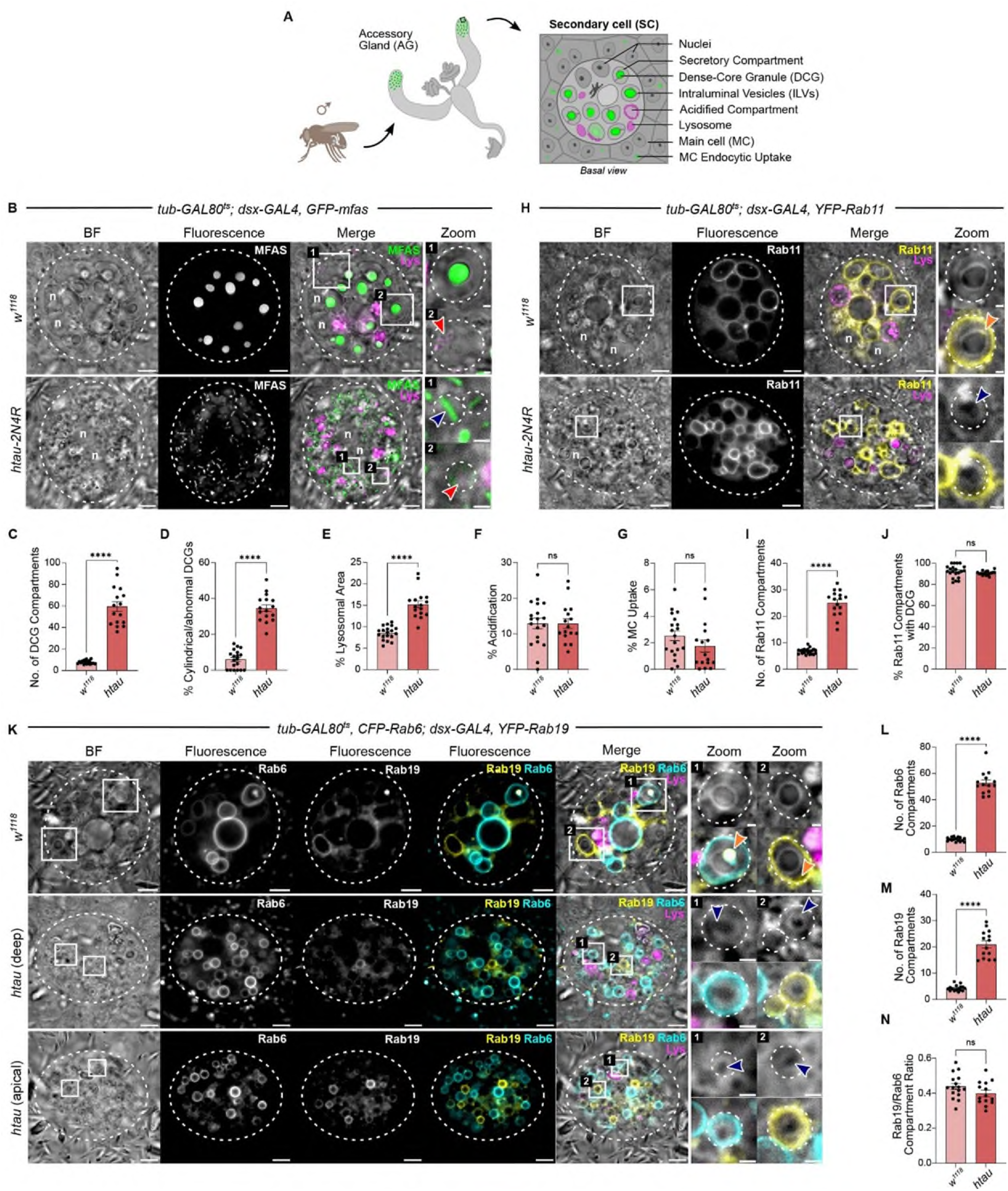
Htau overexpression in SCs induces specific defects in DCG biogenesis and endolysosomal trafficking. **A.** Schematic illustrating male accessory gland system and a single SC, located at the distal tip of a MAG lobe, surrounded by main cells, which can be visualised in a basal view. There are approximately 10 DCG compartments, which normally contain a large (∼3 µm diameter) spherical DCG (marked by GFP-MFAS) and some ILVs that will be secreted as Rab11-exosomes. At most, only one or two DCG compartments are acidified via the SC’s quality control pathway for regulated secretion. **B.** *Ex vivo* wide-field fluorescence images of single SC (marked by dashed circle) in MAGs dissected from males carrying the *GFP-mfas* gene trap fusion and stained with LysoTracker Red. These SCs express no UAS-regulated transgene (top row) or htau(-2N4R). Bright-field view reveals outlines of large secretory and endolysosomal compartments, as well as the two nuclei (n) of these bi-nucleate cells. Zoom images show a DCG compartment (1) with an htau-induced cylindrical DCG (blue arrowhead; also marked in **H** and **K**) and an acidified DCG compartment (2) with internal LysoTracker Red staining (red arrowheads). **C-G.** Bar charts comparing SC DCG compartment number (**C**), % compartments with cylindrical or other abnormal DCGs (**D**), % lysosomal (LysoTracker Red-positive) area (**E**), % of acidified compartments (**F**), and % MC area containing GFP-MFAS (**G**). In **C**, **D** and **F**, a single z-plane where the largest number of DCG compartments are located is analysed. **H.** *Ex vivo* wide-field fluorescence images of single SC (marked by dashed circle) in MAGs dissected from males carrying a *YFP-Rab11* endogenous gene fusion and stained with LysoTracker Red. Note that htau-induced cylindrical DCGs have relatively greater diameter in the absence of GFP-MFAS. Zoom images show a Rab11-positive DCG compartment in the absence and presence of YFP fluorescence with orange arrowheads marking ILVs at the DCG’s periphery in control cells (also marked in **K**). **I, J**. Bar charts comparing SC Rab11-positive compartment number (**I**), and % Rab11-positive compartments with DCG (**J**) for these genotypes. **K.** *Ex vivo* wide-field fluorescence images of single SC (marked by dashed circle) in MAGs dissected from males carrying *CFP-Rab6* and *YFP-Rab19* endogenous gene fusions and stained with LysoTracker Red. Zoom images show a Rab6-positive (1) and Rab19-positive (2) DCG compartment in the absence and presence of fluorescence signal. Apical view of htau-expressing SCs reveals that few Rab19-positive compartments are located peripherally, unlike controls. **L-N.** Bar charts comparing SC Rab6-positive compartment number (**L**), Rab19-positive compartment number (**M**) and ratio of Rab6-to Rab19-positive compartments (**N**) for these genotypes. The images shown are typically a z-stack of 2-3 slices to more fully represent the cellular phenotype. n = nuclei. White squares mark positions of Zoom areas. Scale bars: 5 µm and 1 µm for higher magnification (Zoom) views.

Htau overexpression induced a large increase in the number of DCG compartments, which were reduced in size (Fig. 1B). Indeed, there were too many DCG compartments to score accurately throughout the cell, so as a measure of compartment number, we counted DCG compartments in a central z-plane that included the highest number of compartments in cross-section (Fig. 1C). Notably, about 40% of these compartments contained cylindrical, and not spherical, DCGs with at least one end of the cylinder in close contact with the compartment’s limiting membrane (Figs. 1B and 1D).

Individual lysosomes were also smaller in htau-expressing cells. To assess lysosome content in SCs, we determined their total area (Singh et al., 2025); however, because they were more difficult to distinguish from acidified DCG compartments, we again focused on the same single z-plane. When compared to controls, total lysosomal area in this plane was significantly elevated with htau overexpression (Fig. 1E), indicating increased targeting of DCG compartments for lysosomal degradation. This idea was consistent with the observation that about 10% of the large number of DCG compartments in the same plane appeared to be acidified (Fig. 1F), an intermediate step in the lysosomal targeting process (Singh et al., 2025; Fig. 1A). However, unlike when SCs express Aβ-peptides, endocytosis of SC-secreted GFP-MFAS by neighbouring epithelial main cells (MCs), was not affected by htau overexpression (Fig. 1G).

To assess whether any of these defects might be explained by altered DCG compartment maturation, we analysed htau-overexpressing SCs in flies producing endogenous levels of a YFP-tagged form of Rab11 (Fig. 1H), a recycling endosomal marker, which selectively labels almost all DCG compartments in wild type cells, but not their precursor compartments associated with the *trans*-Golgi network (Wells et al., 2023). When compared to htau-expressing SCs from males harbouring the *GFP-mfas* gene trap, there were less DCG compartments in the *YFP-Rab11* genetic background (Fig. 1I), a finding we have reported previously (Marie et al., 2023). Cylindrical DCGs with greater diameter than those in *GFP-mfas* flies were observed in many of these compartments using bright-field microscopy (Fig. 1H Zoom). Almost all of these compartments were Rab11-positive (Fig. 1J), suggesting maturation to recycling endosomal identity had occurred appropriately. The proportion of DCG compartments labelled by Rab6, a marker for immature DCG compartments that is retained during subsequent early stages of DCG maturation (Wells et al., 2023), was also unaffected by htau expression (Fig. S1A-C).

Previous studies have suggested that Rab19 labels the entire limiting membrane of the two or three most mature DCG compartments in SCs (Wells et al., 2023). To determine whether DCG compartments in htau-expressing SCs mature to this stage, we expressed htau in SCs from flies containing CFP-Rab6 and YFP-Rab19 fusions at the endogenous *Rab* gene loci (Fig. 1K). The ratio of Rab19-to Rab6-positive compartments was unchanged compared to controls (Figs. 1L-N), suggesting that the complete DCG maturation process takes place in these cells. However, almost all of these Rab19-positive compartments were located centrally in the cell, unlike controls (Fig. 1K), consistent with the possibility that they might not be trafficked normally in the presence of htau.

Recently, we showed that Aβ expression in SCs promotes the accumulation of secreted GFP-MFAS not just in neighbouring MCs, but throughout the MAG, without affecting levels of GFP-MFAS secretion (Cording et al., 2026). We assessed GFP-MFAS endocytosis by MCs in the centre of MAGs with SC-specific htau overexpression and found no detectable changes in this or SC secretion (Fig. S1D-I), suggesting that htau expression does not affect this process in the same way.

We conclude that htau expression in SCs disrupts DCG morphology, generating a unique cylindrical DCG structure that appears to maintain association with the DCG compartment’s limiting membrane. This also leads to increased targeting to lysosomes and a build-up of many smaller DCG compartments.

### Htau overexpression disrupts early stages of DCG compartment maturation

Expression of mutant Aβ-peptides in SCs disrupts the earliest stages of DCG protein aggregation during compartment maturation, preventing dissociation of aggregates from the compartment’s limiting membrane and reducing the mobility of these compartments within the cell, potentially via altered interactions with the cytoskeleton (Singh et al., 2025). To assess whether tau overexpression also affects early steps in DCG aggregation, we followed these events in real-time with the GFP-MFAS marker.

Remarkably, even before DCG aggregation was triggered in immature compartments, the cloud of GFP-MFAS inside these compartments was localised to an apple-core-like structure that contacted the limiting membrane at its ends (Fig. 2A). Aggregation foci formed within this cloud, but they could only stabilise in close proximity to the limiting membrane, after which a wave of aggregation travelled through the cloud, creating a cylindrical DCG. Therefore, htau overexpression affects intra-compartmental DCG maturation events, even before protein aggregation is initiated.

**Fig. 2.**
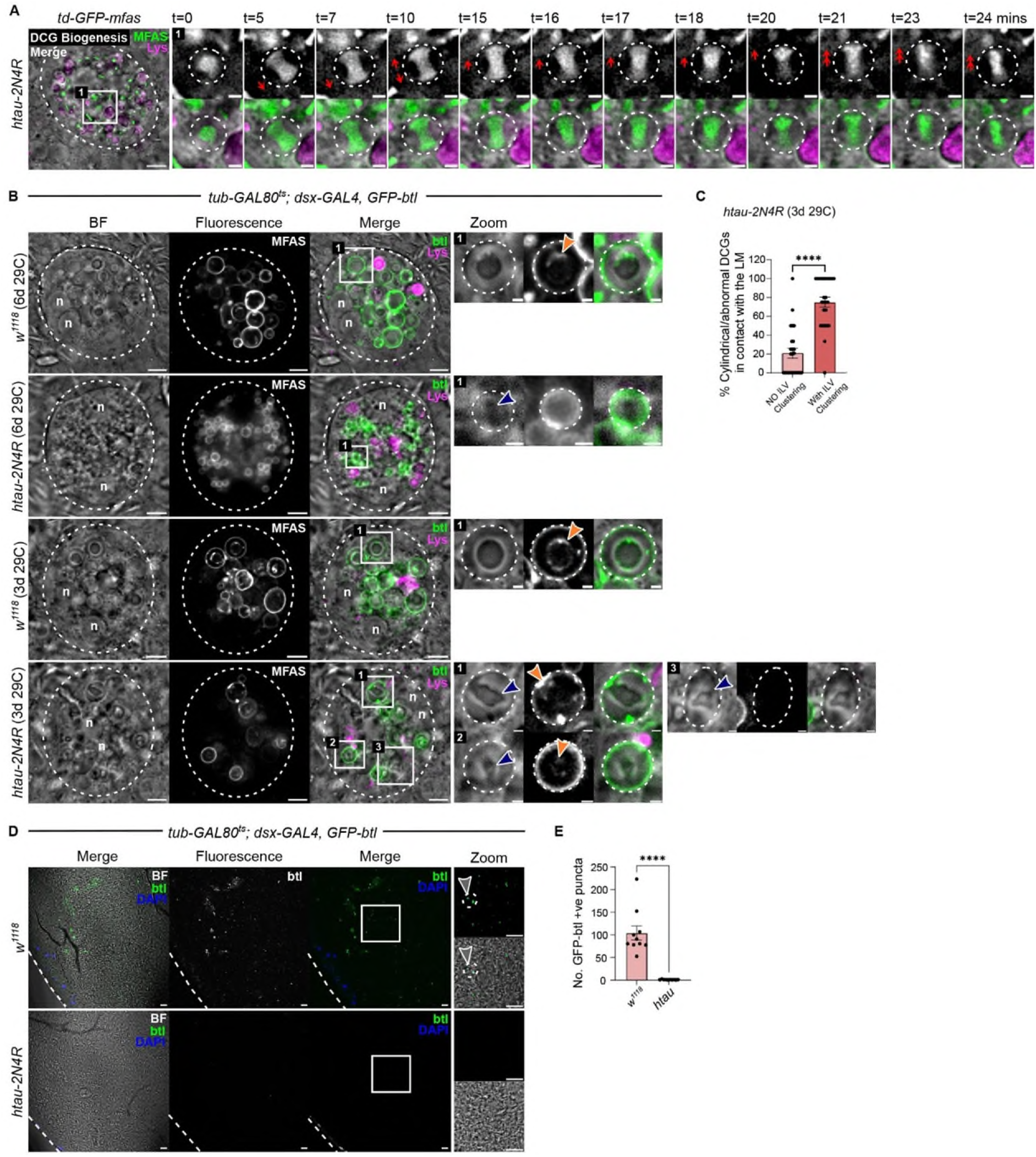
SC-specific htau overexpression modulates early events in exosome biogenesis and DCG protein localisation and aggregation. **A.** Individual frames (in Zoom images) over 24 minutes from an *ex vivo* wide-field fluorescence movie of an early maturing DCG compartment from a single LysoTracker Red-stained SC (marked by dashed circle) in MAGs dissected from males carrying the *GFP-mfas* gene trap fusion and expressing htau from days 4-6 of adulthood. The cloud of non-aggregated GFP-MFAS rotates 90 degrees in the first 5 minutes to reveal an apple-core-like structure. Aggregation events appear to occur sporadically, but then stabilise at one membrane-associated end of the cloud (t = 20), followed by a wave of aggregation. **B**. *Ex vivo* wide-field fluorescence images of single SC in LysoTracker Red-stained MAGs dissected from males expressing the exosome marker Btl-GFP in the absence and presence of htau. Co-expressing htau and Btl-GFP over 6 days induces formation of very small DCG compartments, which cannot be easily scored for cylindrical DCGs (top two rows). Induction of htau and Btl-GFP expression from days 4-6 overcomes this problem. Zoom images show individual DCG compartments. In the absence of htau expression, Btl-GFP-labelled ILVs (orange arrowhead) are found at the periphery of the central DCG. However, in DCG compartments from htau-expressing SCs, which often contain a cylindrical DCG (blue arrowhead), the ILVs are most commonly located at the limiting membrane adjacent to the DCG (1). However, they are sometimes found more centrally in association with the DCG (2), while in other cases, the compartment cannot be scored, because it is unlabelled by Btl-GFP (3), a phenotype that is not observed in controls. **C.** Bar chart showing % of compartments with cylindrical DCGs in htau-expressing SCs that have Btl-GFP-positive ILVs clustered at the DCG’s point of contact with the limiting membrane. **D.** Images from lumen in central region of MAGs in which SCs express Btl-GFP in the absence and presence of htau. **E.** Bar chart showing number of Btl-GFP-positive puncta in the MAG lumen for these two genotypes. n = nuclei. White squares mark positions of Zoom areas. Scale bars: 5 µm and 1 µm for higher magnification (Zoom) views.

We conclude that the effects of Aβ and tau converge on some of the earliest post-Golgi events in regulated secretion, and ultimately lead to increased targeting of DCG compartments to lysosomes, a trafficking pathway that potentially increases Aβ generation (Verma et al., 2026). However, unlike defective DCG compartments in Aβ-expressing SCs (Singh et al., 2025), htau does not seem to restrict rotational motility of the DCG compartment, although many compartments appear to remain at the same 3D-coordinates within the cell (Fig. 2A), suggesting some defects in intracellular trafficking.

### Htau overexpression also affects ILV formation in SC DCG compartments and suppresses subsequent Rab11-exosome secretion

DCG aggregation in SCs initiates at the limiting membrane of maturing DCG compartments and is followed by dissociation of the resulting aggregates in a process that involves cleavage of *Drosophila* APP, APP-like (APPL); this function can be substituted by human APP (Singh et al., 2025). Initial aggregation events are often in close proximity to peripheral clusters of ILVs in each compartment. As the large central DCG assembles, these ILVs form chains that connect the spherical aggregate to the limiting membrane (Wells et al., 2023). Previous studies suggest that suppressing genes required for ILV biogenesis in SC DCG compartments can disrupt DCG assembly (Marie et al., 2023). Furthermore, large DCG assembly also involves a non-glycolytic, extra-vesicular activity of the enzyme, Glyceraldehyde-3-phosphate dehydrogenase (GAPDH), which clusters ILVs, exosomes and EVs (Dar et al., 2021). We therefore tested whether ILV biogenesis or clustering in maturing DCG compartments is modulated by htau overexpression, using an overexpressed GFP-tagged form of the *Drosophila* FGF receptor Breathless (Btl), which labels SC Rab11-ILVs and secreted Rab11-exosomes (Fan et al., 2020).

Btl-GFP overexpression induces an increased number of DCG compartments of reduced size (Fan et al., 2020). Co-expression of htau and Btl-GFP for six days in adult SCs produced many DCG compartments that were too small to accurately identify ILVs and cylindrical DCGs (Fig. 2B). We therefore changed the induction protocol, ageing adults at 25°C for three days before activating transgene expression at 29°C for a further three days. Under these conditions, many DCG compartments still contained cylindrical DCGs, and in nearly 80% of these, Btl-GFP accumulated adjacent to the limiting membrane at one of the contact sites with the DCG (Figs. 2B and 2C).

To assess whether aberrant htau-induced attachment of ILVs to the limiting membrane of DCG compartments might suppress secretion of these vesicles as Rab11-exosomes, we counted Btl-GFP puncta in the lumen of the accessory gland with and without expression of htau, a standard assay for Rab11-exosome secretion in this system (Fan et al., 2020; Dar et al., 2021; Marie et al., 2023). Despite the fact that GFP-MFAS secretion appears unaffected by htau expression (Figs. S1D and S1I), Rab11-exosome secretion was very strongly suppressed (Figs. 2D and 2E).

Therefore, htau overexpression interferes with DCG biogenesis in SCs, generating a unique cylindrical DCG phenotype. It disrupts dissociation of ILVs and DCG aggregates from the secretory compartment’s limiting membrane, effects that have previously been reported when Aβ is expressed in these cells. However, unlike Aβ, htau suppresses secretion of Rab11-exosomes, though not DCG cargos.

### *cofilin* knockdown phenocopies htau-induced phenotypes in secondary cells

We reasoned that the effect of htau overexpression on ILV and DCG biogenesis likely involves pathological interactions with cytoskeletal regulators normally expressed in SCs. The actin cytoskeleton has been linked to some forms of endosomal ILV biogenesis via its interactions with the adaptor and scaffolding protein Syntenin-1 (Lee et al., 2023). Although there is no Syntenin orthologue in *Drosophila*, orthologues of other proteins involved in Syntenin-1’s regulation of ILV production, namely Syndecan and ALIX, are present in flies. Since multiple regulators of the actin cytoskeleton have been implicated in AD (Haseena et al., 2026; Table S1), we investigated whether knockdown of *Drosophila* orthologues of these molecules could also induce a similar phenotype to htau overexpression in SCs (Table S1).

Although suppressing expression of several of these genes disrupted DCG compartments, knockdown of *cofilin* (*tsr* in *Drosophila*), which encodes an actin-severing protein (Huang et al., 2006), uniquely generated SCs in which cylindrical DCGs are formed (Figs. 3A and 3C; Table S1). The phenotype observed in different SCs was somewhat variable: in many SCs, most compartments contained cylindrical DCGs, while the phenotype was more mixed in other cells. In about 20% of cells, no cylindrical DCGs were generated. However, in all cases, unlike controls, all DCG compartments did not occupy peripheral locations in the cell, suggesting that their overall intracellular trafficking was disrupted by the cytoskeletal changes induced.

**Fig. 3.**
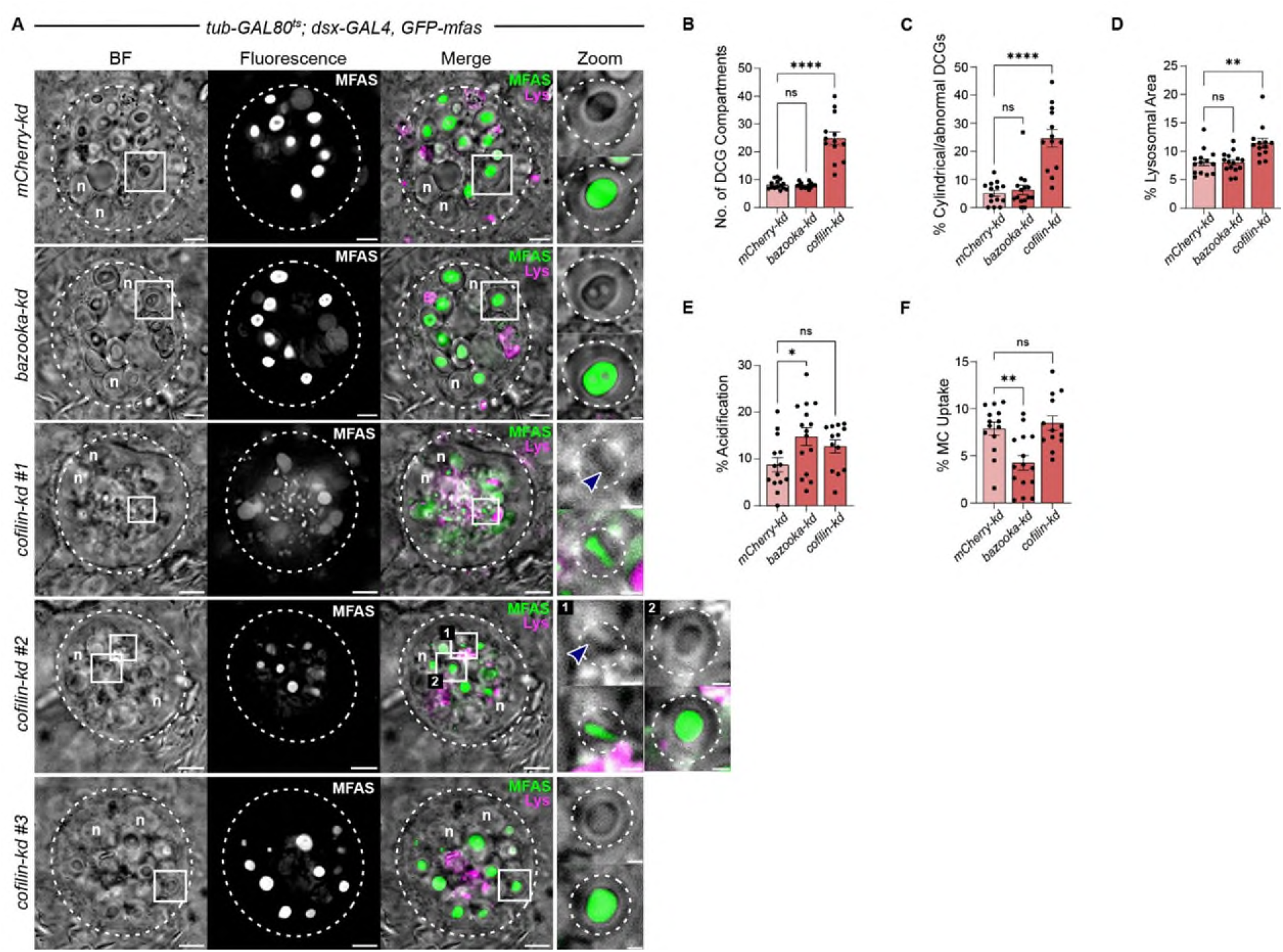
*cofilin* knockdown induces a cylindrical DCG phenotype and multiple trafficking defects. **A.** *Ex vivo* wide-field fluorescence images of single SC (marked by dashed circle) in MAGs dissected from males carrying the *GFP-mfas* gene trap fusion and stained with LysoTracker Red. These SCs express RNAis targeting *mCherry*, *baz* or *cofilin* (bottom three rows). Zoom images show a DCG compartment in the absence and presence of GFP fluorescence. The effects of *cofilin* knockdown vary: in the most common phenotype (#1), many DCGs are cylindrical, while a few cells exclusively contain spherical DCGs (#3) and others (#2) have both cylindrical (1) and spherical (2) DCGs, explaining the broad spread of data in **B** and **C**. However, all cells lack DCG compartments at their periphery. **B-F.** Bar charts comparing SC DCG compartment number (**B**), % compartments with cylindrical or other abnormal DCGs (**C**), % lysosomal (LysoTracker Red-positive) area (**D**), % of acidified compartments (**E**), and % MC area containing GFP-MFAS (**F**). n = nuclei. White squares mark positions of Zoom areas. Scale bars: 5 µm and 1 µm for higher magnification (Zoom) views.

As well as forming cylindrical DCGs, in most *cofilin* knockdown SCs, DCG compartment number was elevated (Fig. 3B) and there appeared to be a modest increase in lysosomal area (Fig. 3D) with no significant effect on acidification (Fig. 3E) and transfer of GFP-MFAS to MCs (Fig. 3F), again mirroring the effects of htau overexpression. By contrast, knockdown of a control gene *bazooka* (*baz*), which encodes the *Drosophila* orthologue of mammalian Par-3, a key protein involved in organising linkage between microfilaments and adherens junctions (Harris and Peifer, 2005), had no obvious effect on DCG compartments, when compared to the effects of an RNAi targeting *mCherry* (Fig. 3). However, for *baz* knockdown, there was evidence of a slight increase in acidification of DCG compartments and reduced build-up of GFP-MFAS in adjacent endocytosing MCs (Figs. 3E and 3F respectively), supporting the idea that the actin cytoskeleton plays a multifaceted role in DCG compartment maturation and release.

Overall, we conclude that depleting levels of Cofilin induces a DCG phenotype that mirrors the effects of htau overexpression, though it also appears to impact other processes controlling DCG compartment intracellular trafficking. This suggests that htau’s effects on DCG biogenesis involves a subcellularly localised reduction in microfilament modelling, consistent with the previous finding that stabilisation of the actin cytoskeleton plays an important role in tau-induced neurodegeneration (Fulga et al., 2007).

### Overexpression of Cofilin partially suppresses Tau-induced DCG phenotypes

To test whether htau’s effects on DCG biogenesis could be explained by changes in the actin cytoskeleton that are modulated by Cofilin, we investigated the impact of overexpressing *Drosophila* Cofilin in SCs in the absence and presence of htau. Overexpressing wild type Cofilin alone in SCs had no obvious effect on any aspect of regulated secretion and quality control, although there did appear to be an increased level of GFP-MFAS accumulation in surrounding MCs in this genetic background (Fig. 4A-F).

**Fig. 4.**
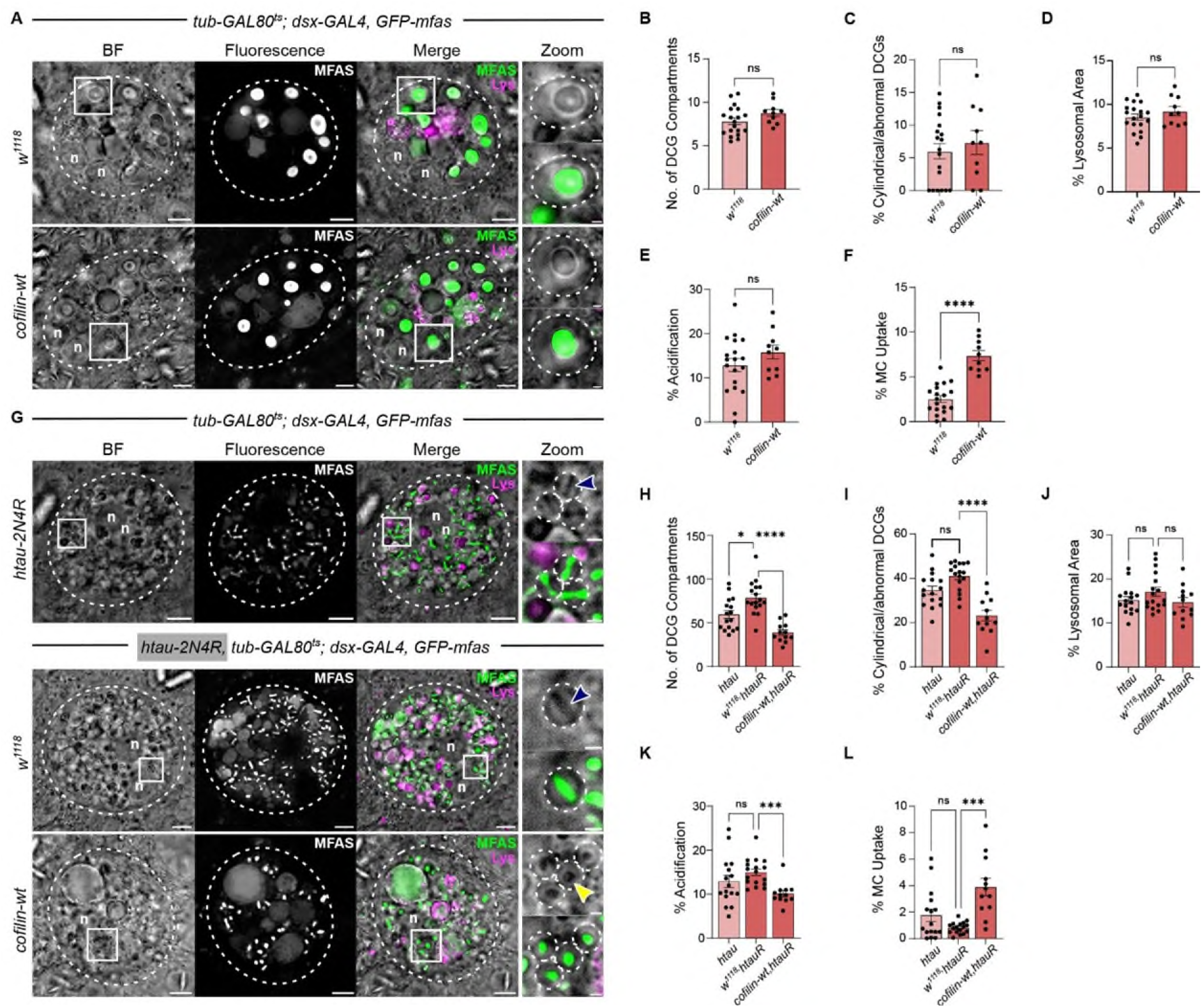
Cofilin co-expression partially suppresses the cylindrical DCG phenotype induced by htau overexpression. **A.** *Ex vivo* wide-field fluorescence images of single SC (marked by dashed circle) in MAGs dissected from males carrying the *GFP-mfas* gene trap fusion in the absence or presence of *UAS-Cofilin* and stained with LysoTracker Red. Zoom images show a DCG compartment in the absence and presence of fluorescence signal. **B-F.** Bar charts comparing SC DCG compartment number (**B**), % compartments with cylindrical or other abnormal DCGs (**C**), % lysosomal (LysoTracker Red-positive) area (**D**), % of acidified compartments (**E**), and % MC area containing GFP-MFAS (**F**). **G.** *Ex vivo* wide-field fluorescence images of single SC in LysoTracker Red-stained MAGs dissected from males carrying the *GFP-mfas* gene trap fusion, and expressing htau in the absence and presence of Cofilin. The MAG in the top row is generated from a genetic cross with the *UAS-htau* line. In the bottom two rows, a recombinant line including the *UAS-htau* transgene has been crossed to control or *UAS-cofilin* flies. Note the htau-induced cylindrical DCG phenotype (blue arrowheads) is suppressed by Cofilin co-expression, producing a spherical DCG (yellow arrowhead). **H-L.** Bar charts comparing same variables as in **B-F** respectively. n = nuclei. White squares mark positions of Zoom areas. Scale bars: 5 µm and 1 µm for higher magnification (Zoom) views.

To determine the effects of Cofilin in an htau overexpression background, we generated a recombinant line in which htau could be inducibly expressed under dsx-GAL4 control in the presence of GFP-MFAS. This line behaved similarly to htau overexpression males generated by a simple cross (Fig. 4G-I). Using this line, co-expressing Cofilin with htau suppressed cylindrical DCG formation and the accompanying increase in DCG compartment number (Fig. 4G-I). Supporting the conclusion that Cofilin partially rescues the htau-induced SC phenotypes, there was a small, but significant, reduction in acidified compartments, though no effect on total lysosomal area (Fig. 4J and 4K). There was also enhanced transfer of GFP-MFAS into adjacent MCs (Figs. 4J-L), which mirrored the effects of Cofilin on MC GFP-MFAS accumulation in the absence of htau (Fig. 4F), suggesting that this effect may not be linked to htau’s activity.

In conclusion, our data suggest that htau overexpression and stabilisation of the actin cytoskeleton, which have previously been functionally linked in the induction of neurodegeneration (Fulga et al., 2007), induce similar defects in early DCG biogenesis events. Indeed, Cofilin overexpression, which increases actin remodelling, partially rescues the DCG defects induced by htau, indicating that htau and Cofilin function antagonistically through the actin cytoskeleton to control DCG biogenesis.

### Cofilin overexpression suppresses Aβ-induced defects in a *Drosophila* eye neurodegeneration model

If htau-induced defects in regulated secretion contribute to degeneration in neurons, Cofilin overexpression would be predicted to suppress tau-induced neurodegeneration. Indeed, Fulga et al. (2007) have reported such suppression in assays where Cofilin is co-expressed with htau in both the eye and brain. To confirm this, we expressed htau in the *Drosophila* eye, using the GMR-GAL4 driver, which activates gene expression in differentiating and adult cells of the fly eye. This has been shown to produce a rough eye phenotype (REP) due to morphological changes associated with neurodegeneration (eg. Fulga et al., 2007). Cofilin appeared to induce a mild suppression of the REP, particularly around the centre and anterior part of the eye (Figure S2).

We have previously employed an image analysis pipeline to assess the regularity of eyes in different genetic combinations where human Aβ is expressed in the eye under GMR-GAL4 control (Cording et al., 2026). This assay is based on measuring the angular distribution of the six closest ommatidia for each ommatidium within a circle of fixed size in the middle of the eye (representing about 250 ommatidia per eye; see Materials and Methods). As expected, htau overexpression induced an REP with reduced regularity compared to control eyes (Fig. S2E versus S2A). However, co-expression with Cofilin had no statistically significant effect on this disorganised eye phenotype, or when compared to htau and lacZ co-overexpressing control eyes (Fig. S2E-H). This suggests that our automated assay is unable to detect the suppression effect of Cofilin on the htau-induced morphological defects in the eye, probably because of the high level of disorganisation in these eyes.

Since not only htau, but also Aβ, affect membrane:aggregate interactions in DCG compartments (Singh et al., 2025), we reasoned that Cofilin-modulated stabilisation of the actin cytoskeleton might be involved in Aβ’s effects on DCG assembly. If these changes are important in inducing degeneration, Cofilin expression might suppress Aβ-induced neurodegeneration. To assess this, we expressed wild type Aβ and a mutant form of Aβ, Aβ-Iowa (Grabowski et al., 2001; Singh et al., 2025), associated with early-onset familial AD, in the fly eye, using GMR-GAL4. Both Aβ isoforms produce an REP that can be easily distinguished using the regularity assay (Chouhan et al., 2016; Metsla et al., 2022; Cording et al., 2026; Fig. 5). Co-expression of Cofilin, when compared to non-co-expressing or lacZ-expressing controls, suppressed the defects observed in both Aβ-wt- and Aβ-Iowa-expressing eyes (Fig. 5), while expression of Cofilin alone had no detectable effect on eye morphology (Figs. S2A-D and S3). These findings suggest that Cofilin expression, which is predicted to increase remodelling of the actin cytoskeleton, can partially suppress the neurodegenerative effects of Aβ. Aβ may therefore mediate some of its effects on neuronal survival by stabilising the actin cytoskeleton, potentially by reducing Cofilin activity. Since Cofilin can suppress both Aβ- and tau-induced phenotypes, these findings are consistent with a model in which modulating the dynamics of the microfilament cytoskeleton and misregulation of DCG biogenesis play important roles in linking the pathological functions of these two major players in AD.

**Fig. 5.**
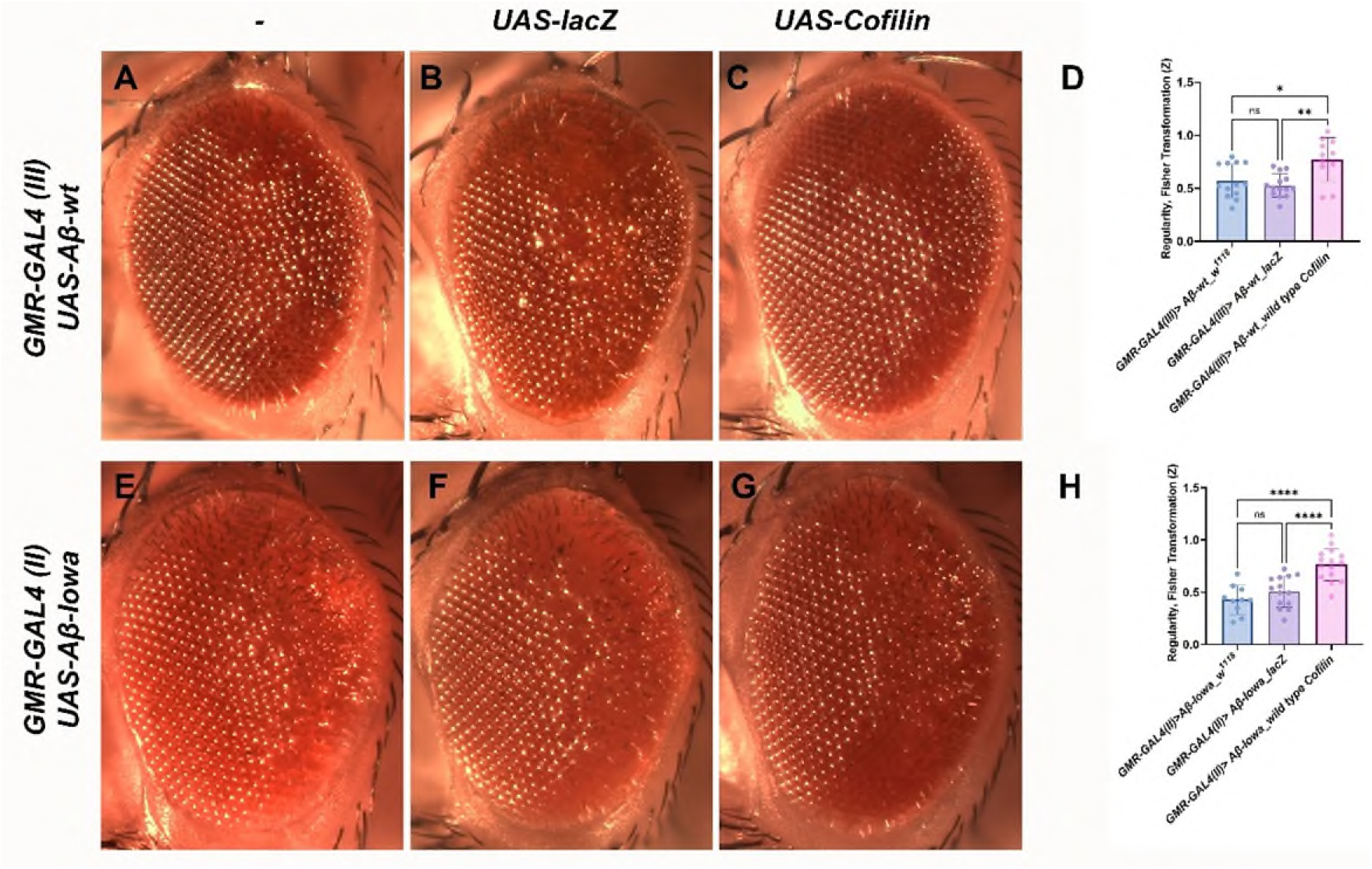
Cofilin co-expression partially suppresses the morphological defects induced by Aβ in the *Drosophila* eye. **A-C.** Stereomicroscopic images of eyes from 2-4-day-old females carrying the third chromosome *GMR-GAL4 (III)* insertion in the presence of *UAS-Aβ-wt* and co-expressing either no other transgene (**A**), *UAS-lacZ* (**B**) or *UAS-Cofilin* (**C**). **D.** Bar chart plotting regularity score (as a Fisher Z transformation) for eyes of these genotypes relative to control *GMR-GAL4 (III)* reference curve. Note partial rescue of disorganisation by Cofilin overexpression relative to both controls. **E-G.** Stereomicroscopic images of eyes from 2-4-day-old females carrying the second chromosome *GMR-GAL4 (II)* insertion in the presence of *UAS-Aβ-Iowa* and co-expressing either no other transgene (**E**), *UAS-lacZ* (**F**) or *UAS-Cofilin* (**G**). **H.** Bar chart plotting regularity score (as a Fisher Z transformation) for eyes of these genotypes relative to control *GMR-GAL4 (II)* reference curve. Cofilin overexpression partially rescues Aβ-Iowa-induced disorganisation relative to both controls. Scale bar: 100 µm.

## Discussion

Despite the well-established association of amyloid plaques and neurofibrillary tangles with AD, the key question of how Aβ and tau might connect and function together to initiate pathology remains unresolved. There is extensive evidence that Aβ disrupts endolysosomal trafficking at early stages of AD (Kimura and Yanagisawa, 2018). Recent studies of DCG compartment biogenesis in SCs have revealed that the quality control pathway in regulated secretion provides a route to generate Aβ physiologically and induce endolysosomal trafficking defects, when DCG aggregation is disrupted and/or Aβ accumulates abnormally (Wells et al., 2023; Singh et al., 2025; Verma et al., 2026). Interestingly, a systems biology analysis of human cellular protein expression and protein:protein interactions within the secretory pathway highlighted defects in secretion and associated protein aggregation, together with downstream effects on endosomal trafficking, as early perturbations in driving AD pathology, supporting this model (Kuo et al., 2021).

Here we show that overexpressing human tau, a manipulation routinely used in *Drosophila* models of AD to induce neurodegeneration, has a highly specific effect on early DCG biogenesis events in SCs (Figs. 1 and 2), which is phenocopied by knockdown of *Drosophila cofilin* (Fig. 3). Increased Cofilin expression not only reduces the DCG phenotype (Fig. 4) and neurodegeneration (Fulga et al., 2007) induced by htau, but also suppresses Aβ-induced defects in an eye neurodegeneration model (Fig. 5). This supports an important role for microfilament dynamics in connecting tau and Aβ to their downstream effects on regulated secretion, endolysosomal trafficking and neuronal survival.

### Overexpression of htau induces defects in early maturation events during regulated secretion

Although htau is primarily thought to operate in the cytosol to regulate the microtubule cytoskeleton, we found that when overexpressed, it induces highly specific defects inside SC secretory compartments, generating a cylindrical DCG without detectably changing the Rab transitions that accompany compartment maturation (Fig. 1). Stable aggregation appears to require priming from the limiting membrane or associated ILVs (Fig. 2), consistent with our previous findings (Singh et al., 2025). However, unlike normal cells, many DCG aggregates do not subsequently appear to dissociate from the limiting membrane, a phenotype also observed following Aβ overexpression. It was difficult to properly characterise DCG compartments that were targeted for lysosomal degradation, because of their small size, but similar to Aβ-expressing SCs, it appeared that more compartments were targeted to lysosomes (Fig. 1).

Aside from the unusual morphology of the defective DCGs induced by htau, there are at least two other differences when compared to Aβ-expressing SCs. First, most compartments where cylindrical DCGs are formed contain ILVs that maintain a close association with both the limiting membrane and the end of the DCG cylinder (Fig. 3), suggesting that tau also affects the pattern of ILV biogenesis and/or organisation in these defective compartments. Second, there is no evidence for propagation of the endolysosomal phenotype produced in SCs to adjacent MCs (Fig. 1G). This may be explained by the observation that htau overexpression blocks Rab11-exosome secretion (Fig. 3), since these exosomes are required for intercellular propagation of endolysosomal defects in the AG system (Cording et al., 2026).

One other surprising finding in htau-expressing SCs is that GFP-MFAS, a major DCG protein with a key role in DCG assembly, is localised in a highly specific apple-core-like structure, even prior to aggregation, suggesting that the process of DCG biogenesis involves early events that concentrate DCG proteins within the compartment lumen. In this regard, it has been proposed that the initial steps in insulin granule assembly in pancreatic β-cells involve recruitment of proinsulin to condensates formed by a Chromogranin B scaffold (Parchure et al., 2022). It will be interesting to test whether any chromogranin-like molecules are involved in DCG biogenesis in *Drosophila* SCs, and whether these disrupt the early concentration of GFP-MFAS in htau-expressing cells.

### Cofilin regulates DCG biogenesis mechanisms modulated by tau

We reasoned that htau was likely to interact with cytoskeletal components in controlling regulated secretion in SCs and focused on the microfilament cytoskeleton, partly because of its known functions in ILV biogenesis (Lee et al., 2023) and proposed roles in AD (Haseena et al., 2026). Indeed, the actin cytoskeleton has been implicated in the trafficking and release of DCG compartments near the plasma membrane in neurons and other secretory cells (Bittins et al., 2009; Wang and Richards, 2017). Furthermore, abnormal accumulation of filamentous actin has been reported to be induced by htau in fly and mouse models, and to play a role in neurodegeneration that is exacerbated by Aβ co-expression (Fulga et al., 2007).

Despite testing the functions of multiple genes in SC DCG biogenesis (Redhai et al., 2016; Fan et al., 2020; Dar et al., 2021; Marie et al., 2023; Wells et al., 2023; Singh et al., 2025), only *cofilin* knockdown phenocopies the DCG defects produced by htau overexpression (Table S1 and Fig. 4), strongly suggesting an important functional link between these molecules. The finding that Cofilin co-expression suppresses the effects of htau overexpression (Fig. 5) further supports this idea, although we cannot eliminate that htau and Cofilin function in parallel pathways that antagonistically control DCG biogenesis. Examples are now emerging of cross-talk between microtubules and microfilaments in cytoskeletal assembly processes (Chou et al., 2026), which may explain the htau/actin link. Alternatively, tau is reported to directly interact with actin, providing a potential dynamic bridge between microtubules and microfilaments (He et al., 2009; Elie et al., 2015).

There are many reports implicating Cofilin in AD. Active Cofilin and actin assemble into pathological rod-shaped bundles, which are more abundant in AD patients and correlate with tau pathology (Rahman et al., 2014), though their link to neurodegeneration remains unclear (Bamburg et al., 2021). Other studies have suggested that Cofilin inactivation can play an important role in AD (reviewed in Kang and Woo, 2019; Paciello et al., 2025), with Aβ oligomers, for example, suppressing Cofilin’s actin-severing functions (Gu et al., 2014; Rush et al., 2018). Importantly, Cofilin’s activity is exquisitely regulated in all cell types (Ohashi, 2015; Bamburg et al., 2021), and overexpression of the wild type protein is unlikely to circumvent this regulation, which probably explains the very mild phenotypes induced by this manipulation in the absence of htau in our experiments (Figs. 4 and 5).

Overall, our data suggest that the effects of overexpressed htau on DCG biogenesis involve stabilisation of the actin cytoskeleton, which can be reversed by Cofilin overexpression. Although Cofilin can induce pathological rod-shaped Cofilin/actin bundles in certain scenarios (Bamburg et al., 2021), the previously reported inhibition of htau-induced filamentous actin accumulation by overexpressed *Drosophila* Cofilin in fly neural tissue (Fulga et al., 2007) likely explains why Cofilin acts as a suppressor in our assays. However, the additional impact of *cofilin* knockdown on DCG compartment trafficking (Fig. 3) suggests that htau’s effects may be subcellularly localised. Presumably, a more dynamic extra-compartmental cytoskeleton is required to permit the normal biogenesis of ILVs, condensation of DCG components and dissociation of DCG aggregates from membranes during compartment maturation (Fig. 6). Since endolysosomal defects in htau-overexpressing SCs are only modestly suppressed by Cofilin, it seems likely that mechanisms other than stabilisation of the actin cytoskeleton play important roles in this aspect of htau-induced secretory pathology.

**Fig. 6.**
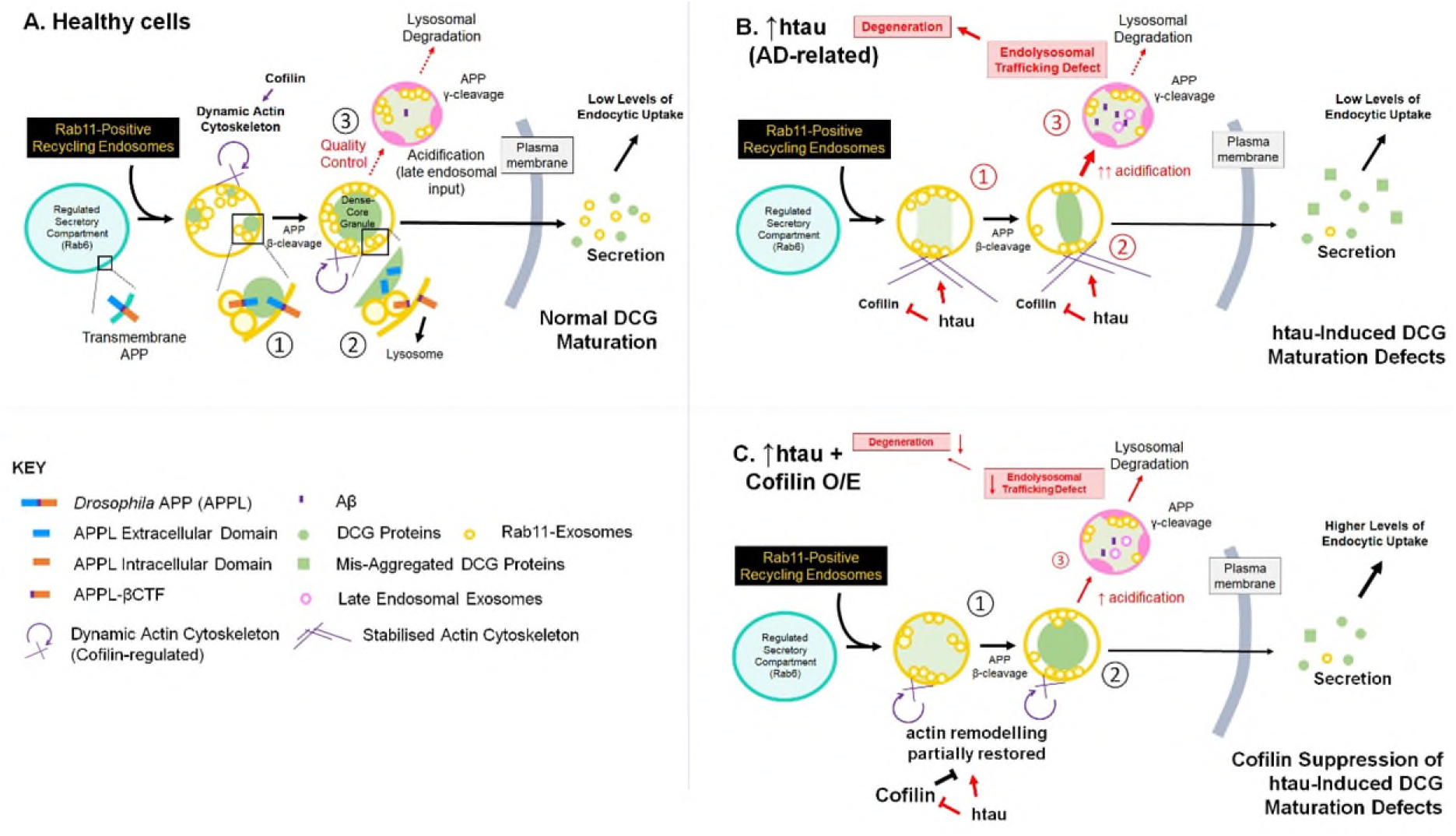
Proposed model to explain Cofilin’s roles in DCG compartment maturation and quality control, and the link to neurodegeneration. Schematic shows the regulated secretory process in healthy cells (**A**), and in htau-expressing cells, in the absence (**B**) or presence (**C**) of overexpressed Cofilin. Regulated secretion involves DCG biogenesis (① and ②), a process requiring a compartmental switch to Rab11-positive recycling endosomal identity and APP processing, followed by quality control (③), and secretion. In healthy cells (**A**), only a small proportion of DCG compartments are targeted for lysosomal degradation, leading to minimal accumulation of Aβ. Normal DCG biogenesis is dependent on Cofilin activity, which maintains a dynamic actin cytoskeleton. We find that if htau is overexpressed (**B**), this disrupts condensation of DCG cargos (red ①) and DCG biogenesis (red ②), at least partly by altering and stabilising the interactions between aggregating DCG proteins and the compartment’s limiting membrane. More compartments are targeted to lysosomes (red ③) and the resulting endolysosomal defects contribute to degeneration. Overexpressing Cofilin in SCs (**C**) partially suppresses this phenotype and neurodegeneration, suggesting a role for actin stabilisation in htau-induced pathology. Note that Cofilin overexpression increases levels of DCG proteins endocytosed by other cells independently of htau via an undetermined mechanism. Aβ expression also inhibits membrane:DCG aggregate dissociation and its neurodegenerative effects in the eye are suppressed by Cofilin, suggesting that the actin cytoskeleton’s effects on DCG compartment maturation may play a critical role in both Aβ- and tau-induced AD pathology.

### Neurodegeneration induced by Aβ is partially suppressed by Cofilin

Although the clinical use of recently developed monoclonal antibodies that disperse amyloid plaques can modestly slow AD progression in a subset of patients (Kim et al., 2025), identifying targets that might suppress initiation of disease remains a major priority. Since Cofilin appears to be a potential player in htau’s effects on DCG biogenesis, it was an obvious candidate to test as a suppressor of Aβ-induced neurodegeneration, given Aβ’s role in DCG formation and compartment maturation. Our findings are consistent with some of the effects of Aβ on regulated secretion involving changes in dynamics of the actin cytoskeleton, providing an important hub that links tau and Aβ effects.

Since Cofilin primarily suppresses htau’s effects on early DCG compartment maturation events, including the defective membrane:aggregate interactions observed, it seems likely that some of Aβ’s effects on these processes are also modulated by Cofilin overexpression. However, although we did not observe strong effects of Cofilin on the endolysosomal trafficking defects induced by htau, further investigation is required to eliminate the possibility that this also plays a role in suppressing Aβ secretory pathology. The microfilament cytoskeleton has been shown to play roles in endolysosomal trafficking that complement the well-established functions of microtubules. Microfilaments are critical in endocytosis, but also control specific steps in trafficking of endosomes to the lysosome (Kjeken et al., 2004; Goebeler et al., 2008; Granger et al., 2014; Cheng et al., 2021), processes that may be partly regulated by Cofilin (Okreglak and Drubin, 2007). Furthermore, reduced Cofilin function has been reported to positively impact APP trafficking and Aβ clearance in rodent models of AD (Liu et al., 2019), suggesting a complex role in other aspects of disease pathology.

The identification of Cofilin as a modulator of Aβ- and htau-induced phenotypes that are of relevance to AD provides a link between these two hallmark-associated proteins (Fig. 6). It suggests that studying regulated secretion and the input of recycling and late endosomal compartments in its quality control can identify and unravel new pathways that might be critical to pathology. Indeed, recently it has been proposed that reduced induction of tau pathology by Aβ may be a key factor in the resilience to dementia observed in some individuals (Aguillar et al., 2025; Lu et al., 2026). In this regard, unbiased proteome localisation studies in human brain tissue have demonstrated co-ordinated reorganisation of endolysosomal proteins associated with early and recycling endosomes, which is more robust in AD patients with dementia versus resilient individuals with high pathology but intact cognition (Jolly et al., 2026). One possibility is that these observations are linked to the Aβ-induced changes in regulated secretion and endosomal trafficking that we have identified (Singh et al., 2025; Verma et al., 2026).

Finally, in sporadic AD, pathology typically emerges over many years and therefore may be due to minor imbalances in causative pathways. Therefore, defining other players in the Aβ- and htau-modulated mechanisms controlling regulated secretion, even if they have very general roles in cell biology, may identify a strategy for restoring this imbalance at the earliest stages of disease, and therefore slowing progression many years before the devastating impact of AD emerge.

## Materials and Methods

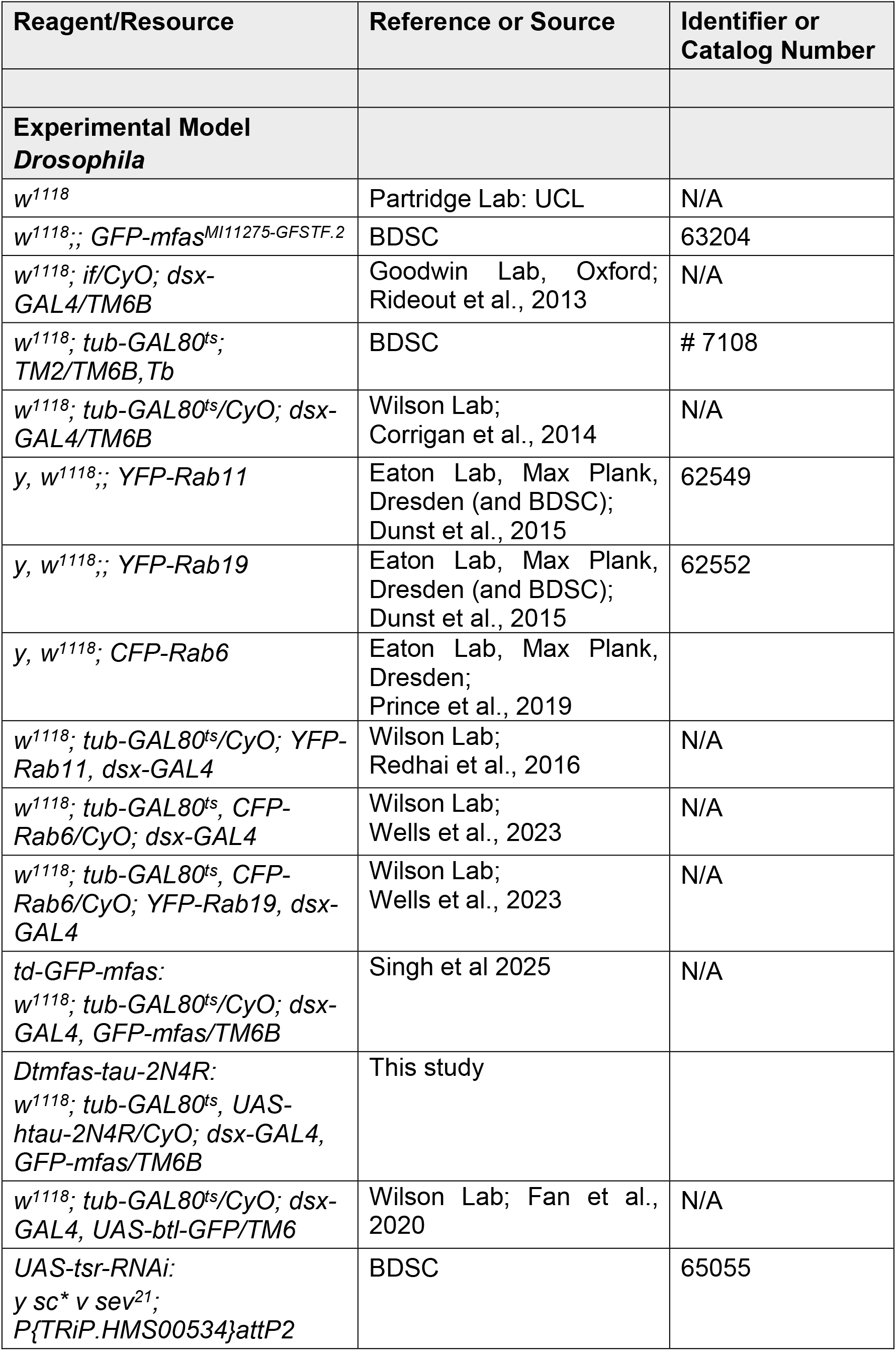
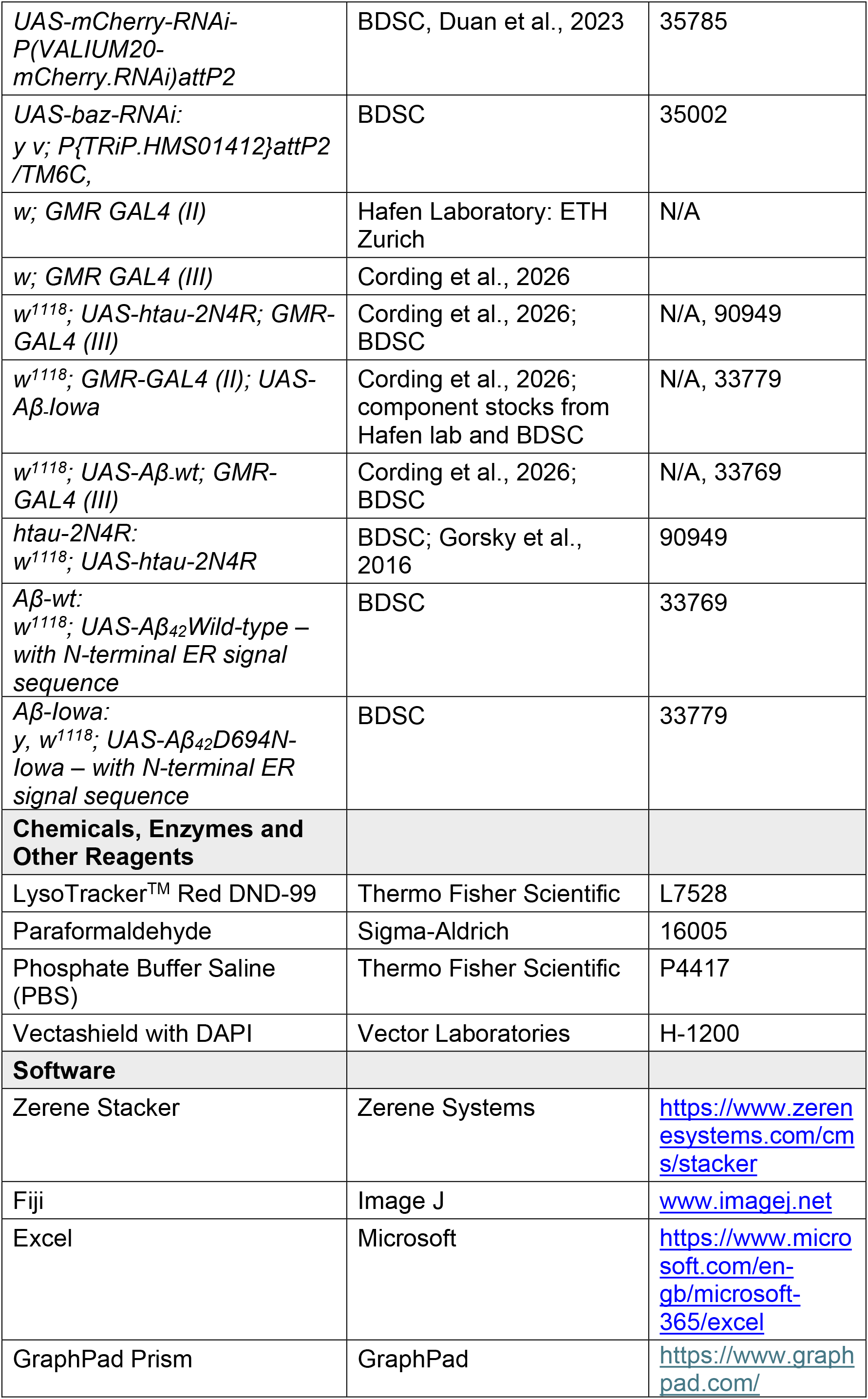

### Fly stocks and husbandry

*Drosophila* strains used in this manuscript are listed in the Reagents table. Most of the transgenic fly lines were obtained from Bloomington *Drosophila* Stock Centre (BDSC) and Vienna *Drosophila* Resource Centre (VDRC). Exceptions were as follows; *dsx-GAL4* (provided by S. Goodwin, Oxford, UK) (Rideout et al, 2010), YFP-*Rab6*, *Rab11*, *Rab19* fusion genes at endogenous *Rab* locus (provided by S. Eaton, Max Plank, Germany) (Dunst et al, 2015; Prince et al., 2019). In cases where lines were generated and/or used in previous studies, this is noted next to the stocks in the list below: *w^1118^* (provided by L.Partridge, UCL,UK) (eg. Singh et al., 2025): *GFP-mfas* gene trap (MI11275 GFSTF.2; BDSC 63204) (Nagarkar-Jaiswal et al, 2015; Singh et al., 2025): *w; tub-GAL80^ts^; dsx-GAL4, GFP-mfas* (Singh et al., 2025): *w; tub-GAL80^ts^; YFP-Rab11, dsx-GAL4* (Fan et al, 2020): *w; tub-GAL80^ts^, CFP-Rab6/CyO; dsx-GAL4* (Wells et al., 2023): *w; tub-GAL80^ts^, CFP-Rab6/CyO; YFP-Rab19, dsx-GAL4* Wells et al., 2023): *UAS-Aβ-wt* (BDSC 33769) (Wu et al, 2017) and *UAS-Aβ-Iowa* (BDSC 33779; Vitruvean) (Chouhan et al, 2016) constructs contain a cleavable ER signal sequence, to direct them to the secretory system (used in Singh et al., 2025): *UAS-mCherry-RNAi* (VALIUM20-mCherry; BDSC 35785) (Duan et al., 2023; Cording et al., 2026): *UAS-tsr-RNAi* (TRiP.HMS00534; BDSC 65055) (Perkins et al., 2015): *UAS-baz-RNAi* (TRiP.HMS01412; 35002): *GMR-GAL4 (II)* (provided by Hafen Laboratory: ETH Zurich) (Freeman, 1996), *GMR-GAL4 (III)* (generated in Wilson lab): *GMR-GAL4 (III*) was combined to create *w; UAS-Aβ-wt; GMR GAL4 (III)* and *w; UAS-htau-2N4R; GMR GAL4 (III)*, and *GMR-GAL4 (II)* used to generate *w; GMR GAL4 (II); UAS-Aβ-Iowa*.

Flies were maintained at 25°C with a 12-hour light/dark cycle on standard cornmeal agar medium [12.5 g agar (F.Gutlind & Co. Ltd), 75 g cornmeal (B. T. P. Drewitt), 93 g glucose (Sigma-Aldrich, #G7021), 31.5 g inactivated yeast (Fermipan Red, Lallemand Baking), 8.6 g potassium sodium tartrate tetrahydrate (Sigma Aldrich, #S2377), 0.7 g calcium chloride dihydrate (Sigma-Aldrich, #21907), and 2.5g nipagin (Sigma-Aldrich, #H5501) dissolved in 12 ml ethanol, per litre]. Flies were transferred onto fresh food every 3-4 days.

For the experiments involving the MAG, female flies carrying the driver line *tub-GAL80^ts^; dsx-GAL4* alone or in combination with *GFP-mfas*, or *YFP-Rab11*, *CFP-Rab6, CFP-Rab6/YFP-Rab19,* or *UAS-Btl-GFP* (in some cases, together with *UAS-htau-2N4R*) were crossed with male flies carrying UAS-transgenes or *w^1118^* (control), permitting a temperature-controlled induction of SC-specific target gene expression.

As in previous studies (Singh et al., 2025), for the experiments involving the MAG and live tissue imaging, virgin male offspring were collected upon eclosion and transferred to 29°C for 6 days to activate post-developmental SC-specific transgene expression. In some experiments with htau overexpression, males were maintained for 3 days at 25°C after eclosion, then switched to 29°C for 3 days to reduce the effect on DCG compartment size and number.

The experiments in which we quantified GFP-MFAS in the main cells at the centre of the MAG involved imaging of fixed tissue (see Cording et al., 2026); virgin males were transferred to 29°C at eclosion to induce htau expression.

For the experiments involving *Drosophila* eye analysis, female flies carrying *UAS-*transgenes or *w^1118^* controls were crossed with male flies carrying *GMR-GAL4* alone or in combination with *UAS-tau-2N4R*, *UAS-Aβ-wt* or *UAS-Aβ-Iowa*, to induce eye-specific expression of target genes (see Cording et al., 2026). These crosses were maintained at 25°C throughout the experiment. Upon eclosion, female flies were separated from males and aged 2-4 days at 25°C before freezing and imaging.

### Preparation of MAGs for imaging

The methodology was similar to that described by Singh et al., 2025 and Cording et al., 2026 for live-cell imaging. Six-day-old adult male virgin flies were anaesthetised using CO_2_, then submerged in ice-cold 1X PBS (Thermo Fisher Scientific). The male reproductive system was dissected out together with the testes, seminal vesicles and ejaculatory bulb, but fat tissues and the gut were removed. The glands were then incubated with 500 nM Lysotracker Red (Thermo Fisher Scientific) in PBS for 2-5min on ice, followed by a wash with ice-cold PBS. The tissue was then mounted between two coverslips (rectangular: coverslip No.1, 22 mm×50mm, Fisher, #1237-3128, and round: coverslip No.1, 13 mm, #49492, VWR) in a drop of 1x PBS, with the set-up held together by a custom-built metal holder. Excess PBS was removed using a filter paper when the glands were ready for imaging, to slightly flatten the samples.

For confocal analysis, micro-dissections were performed in PBS at room temperature and then samples fixed in 4% paraformaldehyde in PBS (Sigma-Aldrich) for 20 min. The glands were washed in 1x PBS for approximately 5 min prior to mounting in a drop of Vectashield with DAPI (Vector Laboratories) on SuperFrost microscope slides (VWR). Samples were held in place with a coverslip (22 mm × 22 mm, 0.13–0.17 mm; Fisher)

### Preparation of flies for eye imaging

Following the protocol in Cording et as., 2026, for light stereomicroscopy of adult eyes, 2-4-day-old females flies were anaesthetised with carbon dioxide and then transferred to 0.6 ml Eppendorf tubes (SLS) and frozen at -20°C for a minimum of 24hrs, with most flies imaged within one week after freezing. To image the eyes, flies were positioned with the central part of the eye as horizontal as possible, facing the objective lens.

### Live-cell imaging

Live-cell imaging was undertaken at room temperature and followed the methodology used in Singh et al., 2025. A Leica Thunder inverted wide-field microscope (Leica) was employed, at 100x magnification (Leica HCX PL FLUOTAR NA 1.3, oil objective) with a K8 sCMOS camera. Four SCs were analysed per MAG (two per lobe) for ≥10 individual virgin males. The images acquired were typically z-stacks spanning a depth of 8–12 μm with a z-distance of 0.2 μm.

Thunder technology (Leica) with small volume computational clearing (SVCC) was applied to enhance contrast and eliminate out-of-focus blur. SC morphology was visualised using FLURO-Bright Field imaging. The LED settings were as follows: GFP, 475 nm excitation at 40% laser intensity; YFP, 510 nm excitation at 42% laser intensity; and RFP, 550nm excitation at 30% laser intensity.

### Fixed-cell imaging

Fixed samples were imaged on a Zeiss LSM980 with Airyscan 2 Super-resolution upright laser scanning confocal microscope equipped with 10x (Zeiss 0.45 NA; dry) and 40x (Zeiss 1.30 NA; oil; Zeiss immersion oil, refractive index 1.518) objectives.

The two lobes of the MAG were imaged for ≥10 individual virgin males. High resolution images of the MAG’s main cells and lumen were acquired from the central region of each lobe employing the 40x objective with 1x zoom on the ZEN bluesuite Software (Zeiss), using GFP, 488 nm excitation at 2% laser intensity and DAPI, 345 nm excitation at 2% laser intensity.

For MC images, the epithelial layer was identified using the DAPI nuclear marker, present in the Vectashield mounting medium. One image was taken through the middle of these nuclei for each lobe for ≥10 individual virgin males.

For lumen images of Btl-GFP, a stack of three z-sections separated by 2 µm was performed that was centred around the middle of the MAG lumen at the centre of both MAG lobes, and this was repeated for at least 10 individual virgin males.

### Eye imaging

This followed the procedure described in Cording et al., 2026.

Brightfield images for qualitative analysis were illuminated by two external LEDs directed either side of the eye.

The GFP excitation wavelength and GFP filter on the camera was used to produce images for quantitative analysis, because the autofluorescence detected produced better contrast for individual ommatidia. Images were captured manually in multiple z-planes using a Leica model MSV269 stereomicroscope with CHROMYX HD camera attachment, at 8X magnification. Between 10-20 consecutive images of each eye were collected, which were then stacked using Zerene Stacker (Zerene Systems, Richland, WA) to produce a composite image of each eye.

### Analysis and parameters

The analysis of SCs and MCs in MAGs employed the approaches previously used by Singh et al., 2025 and Cording et al., 2026. For analysis of SCs in each gland, typically two cells were analysed from each of the two lobes and the mean value calculated per gland for a total of ten or more individuals.

### DCG phenotypes and number of mature compartments

Deconvolved images were analysed using Fiji/ImageJ. Channels were merged to create composite images and the numbers of intact DCGs marked by GFP-MFAS and DCG-containing compartments were quantified manually using the brightfield channel. In analyses where genotypes producing large numbers of DCG compartments were under analysis, counts were derived from the single central z-plane that included the highest number of DCG compartments in cross-section. Cylindrical DCGs, as well as other abnormal cores, which included mini-cores or deformed cores, were then manually counted. At least one end of cylindrical DCGs was in close contact with the limiting membrane of the compartment. Like other misshapen DCGs, the ratio of the lengths of the longest and shortest DCG axes was >1.4. The mini-core phenotype was defined by the presence of at least three small cores of diameter ≥ 0.5 µm.

The percentage of cylindrical/mini-/deformed cores was calculated relative to the total number of DCGs in non-acidic compartments per SC.

To quantify compartments marked by the YFP-Rab11 and CFP-Rab6 fusion proteins, fluorescently labelled compartments were manually examined using single z-planes for the fluorescence channel (Fan et al., 2020; Wells et al., 2023). Because of asymmetric distribution of compartments labelled by different Rabs, all compartments were counted in the analysis of CFP Rab6/YFP-Rab19 co-labelled SCs.

We did not undertake a quantitative analysis of Btl-GFP in all SCs, but for htau-expressing cells, we identified compartments containing cylindrical DCGs in the plane with the most DCG compartments and then scored the proportion of these compartments that exhibited concentrations of Btl-GFP-labelled ILVs at contact points between the cylinder and the compartment’s limiting membrane.

### Acidification of secretory compartments

Mature (rounded) DCG compartments that are associated with acidic structures and potentially undergoing lysosomal clearance (the DCG acidification phenotype; Singh et al., 2025) were scored as acidified compartments. These compartments manifest different phenotypes depending on the stage of acidification. Some have a single peripheral lysosomal structure with a slightly diffuse GFP-positive DCG that has maintained its shape, while others have completely diffuse GFP-MFAS and no obvious DCG structure in brightfield. Some acidified compartments have multiple acidic structures arranged in an arc around the compartment boundary, with or without diffuse GFP-MFAS, while others are covered by many acidified domains, but still retain their spherical shape. The percentage of acidified compartments per SC was determined as the proportion of the total number of both non-acidic DCG-containing compartments and acidified compartments that were acidified in each SC.

To calculate the lysosomal area, a freehand tool on Fiji was employed to outline and measure the area of the SC. The threshold and analyse particles tools were utilised to determine the LysoTracker Red-positive area in the complete projection of the RFP channel. The ratio of LysoTracker Red-positive area to total SC area was calculated for each SC and expressed as % lysosomal area.

### *Ex vivo* analysis of GFP-MFAS-containing main cells

For live images of the distal tip of the MAG, which included one SC and parts of surrounding MCs, the Rectangle tool (700 x 700 pixels) on Fiji was used to determine the area of the field of view, and the freehand tool was employed to outline and measure the area of the SC. Subsequently, the threshold and analyse particles tools were applied to measure the area covered by GFP-MFAS in the surrounding MCs using the complete projection of the GFP channel. The percentage MC area containing GFP-MFAS was calculated within the field of view, having excluded the SC area.

### Analysis of GFP-MFAS-containing main cells in fixed tissue

Using a grid tool on ImageJ, a grid of 25 squares was superimposed over the MC image for the GFP-MFAS channel. Each square was 100 x 100 pixels. In most cases, central squares 7,8,9 were analysed and the mean calculated. In cases where the gland was not flat in these areas, a total of three squares from the central block of nine squares were selected and analysed instead. This was repeated for both lobes of each MAG, and a mean calculated, with 10 individual glands scored for each experimental condition. Analysis was performed using a particle tool, where the squares were thresholded with pixel threshold value set to ≥ 25 (255 maximum) at a consistent exposure, and then the following values were calculated inside each square: number of GFP-MFAS puncta, mean size of each punctum, total area of GFP fluorescence, % area covered by GFP fluorescence, mean intensity of GFP-positive pixels, integrated density (% area of fluorescence x mean fluorescence signal intensity). We also calculated % GFP-positive area/number of puncta to determine % GFP-positive area per punctum, a measure of the size of each structure containing abnormally aggregated material.

### Analysis of GFP-MFAS in MAG lumen using fixed tissue

Using the rectangle tool on ImageJ and a specific confocal image, one square (200 x 200 pixels) was marked in the central luminal region of the middle of each lobe, and a mean GFP-MFAS intensity per lobe was calculated using ImageJ mean intensity function. This was then repeated for the other lobe. The two mean intensity values per animal were averaged again to create a mean intensity value for each animal. This was repeated for 10 male virgin flies.

### Analysis of Btl-GFP puncta in MAG lumen using fixed tissue

For each lobe, using the rectangle tool on ImageJ, one square (200 × 200 pixels) was marked in the middle of each image of the central plane in the lumen. Analysis was performed using a particle tool, where the squares were thresholded with pixel threshold value set to ≥ 20 (255 maximum) at a consistent exposure. The particle function on ImageJ was used to count the number of puncta within this square. This was repeated for the equivalent square positioned 2 µm above and 2 µm below this central plane. These counts were then averaged resulting in a mean value. This was repeated for the other lobe, and the mean of the counts for the two lobes calculated. This was repeated for 10 male virgin flies and the overall mean used as a measure of exosome secretion.

### Image processing and preparation for figures

Live-cell images for this study were prepared using deconvoluted stacks collected from the Leica Thunder microscope. In all resulting figures, images were a composite typically of 2-3 z-slices, chosen for their optimal representation of SC morphology. The fluorescence intensity that best captured the phenotypes of interest was selected and utilised for image processing across all the live-cell data. Consistent projection of slices and settings were applied to generate images for analysis for each genotype shown in the figures. All images were cropped to identical dimensions and focused primarily on SC details.

For fixed-gland images of MCs, a single z-plane through the centre of MC nuclei was acquired using the LSM980 confocal microscope. To ensure consistency in fluorescent intensity quantification, the GFP gain settings were kept constant across all genotypes

### Quantification and statistical analysis

For comparing multiple experimental genotypes with the control, we applied the non-parametric Kruskal–Wallis test followed by Dunn’s multiple comparisons post hoc test. When comparing two groups, a non-parametric Mann–Whitney test was used. These statistical analyses were performed on GraphPad Prism. All graphs displayed in the figures show the mean value for each genotype and include error bars representing the standard deviation (SD). Each experiment was independently repeated at least three times with similar results.

### *Drosophila* eye analysis

#### Image processing

This followed the protocol described in Cording et al., 2026. Following stacking using Zerene Stacker, composite images of each eye were processed using FIJI, with an automated set-up (Cording et al., 2026). In brief, an elliptical area of 576 x 624 pixels was selected at the centre of the eye image. The trainable Weka Segmentation ImageJ plugin (Arganda-Carreras et al., 2017) was trained on a series of reference images of the fly eye to detect individual ommatidia and define boundaries between them, in ‘normal’ and disordered eyes. The resulting segmented probability map, converted to 8-bit, was made binary and the watershed transformation applied, to allow identification of individual ommatidia. After processing, the inbuilt FIJI Analyse Particles function generated XY coordinates of ommatidia, which were exported to Excel.

#### Data analysis

For each ommatidium in the selected area, the angle to the nearest six ommatidia was calculated. These angles were summed for each eye of a specific genotype (approximately 250 ommatidia per eye). For the reference curve, the angles for approximately 15 females of the control genotype were summed after standardising the curve by setting the value of the most commonly occurring angle to zero for each eye. The curves for individual eyes of all other genotypes were then compared to this reference curve, again after setting the value of the most commonly occurring angle to zero, and the Pearson Correlation coefficient calculated. The Pearson correlation coefficient relative to the reference curve was reduced for disorganised eyes under test; these values were then subjected to the Fisher transformation for each eye, producing normally distributed data for statistical analysis. The transformed Z-values were exported to GraphPad Prism, where comparisons were made using one-way Brown-Forsythe and Welch ANOVA tests, followed by unpaired t-tests with Welch’s correction for multiple comparisons.

## Acknowledgements

We are grateful to all the staff at the Micron Bioimaging Facility for their support. We thank Suzanne Eaton, Stephen Goodwin, Elodie Prince, Francois Karch and Linda Partridge, as well as the Bloomington and Vienna *Drosophila* Stock Centres for *Drosophila* stocks. We acknowledge the support of the BBSRC (BB/R004862/1, BB/W00707X/1, BB/W015455/1) and Cancer Research UK (C19591/A19076). For the purpose of Open Access, the author has applied a CC BY public copyright licence to any Author Accepted Manuscript (AAM) version arising from this submission.

## Supplementary Figures

**Fig. S1.**
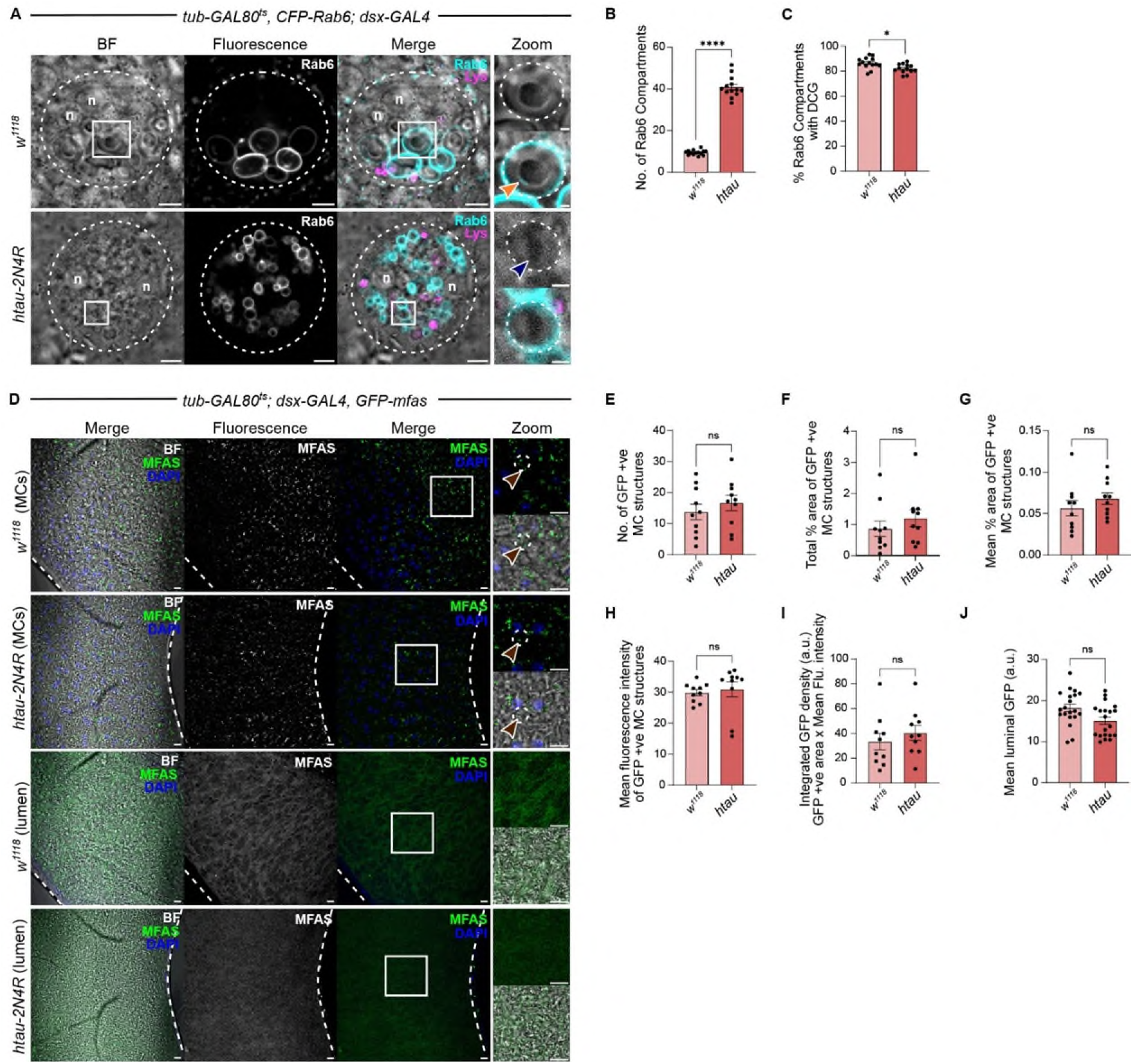
SC-specific htau expression affects neither changes in Rab identity in maturing DCG compartments nor DCG protein secretion. **A.** *Ex vivo* wide-field fluorescence images of single SC (marked by dashed circle) in MAGs dissected from males carrying a *CFP-Rab6* endogenous gene fusion and stained with LysoTracker Red. These SCs express no UAS-regulated transgene (top row) or htau(-2N4R) (bottom). Note that htau-induced cylindrical DCGs (blue arrowhead) have relatively greater diameter in the absence of GFP-MFAS. Zoom bright-field images show a Rab6-positive DCG compartment with and without CFP fluorescence signal. Orange arrowhead highlights CFP-Rab6-labelled ILVs, which are frequently seen in control DCG compartments (Wells et al., 2023). **B, C**. Bar charts comparing SC Rab6-positive compartment number (**B**), and % DCG compartments that are Rab6-positive (**C**), assessed using a single z-plane where the largest number of DCG compartments are located. **D.** Confocal images of the central part of lobes from fixed MAGs dissected from males carrying the *GFP-mfas* gene trap fusion and stained with DAPI to highlight MC nuclei. Top two rows are sections through MC epithelial layer, marked by DAPI-stained nuclei, showing endocytic uptake of GFP-MFAS (aubergine arrowheads). Bottom two rows are sections through middle of MAG lumen showing secreted GFP-MFAS. **E-J.** Bar charts comparing number of GFP-positive structures in MC epithelial layer (**E**), total % area of GFP-MFAS fluorescence in MC layer (**F**), mean % area of individual GFP-MFAS-labelled MC compartments (**G**), mean fluorescence intensity of GFP-positive structures (**H**), integrated density (area multiplied by mean GFP fluorescence intensity in MCs) (**I**), and mean GFP-MFAS intensity in MAG lumen (**J**), none of which are significantly altered by htau expression. SC images are a z-stack of 2-3 slices. n = nuclei. White squares mark positions of Zoom areas. Scale bars: 5 µm and 1 µm for higher magnification (Zoom) views in **A**; 10 µm and 5 µm for higher magnification in **D**.

**Fig. S2.**
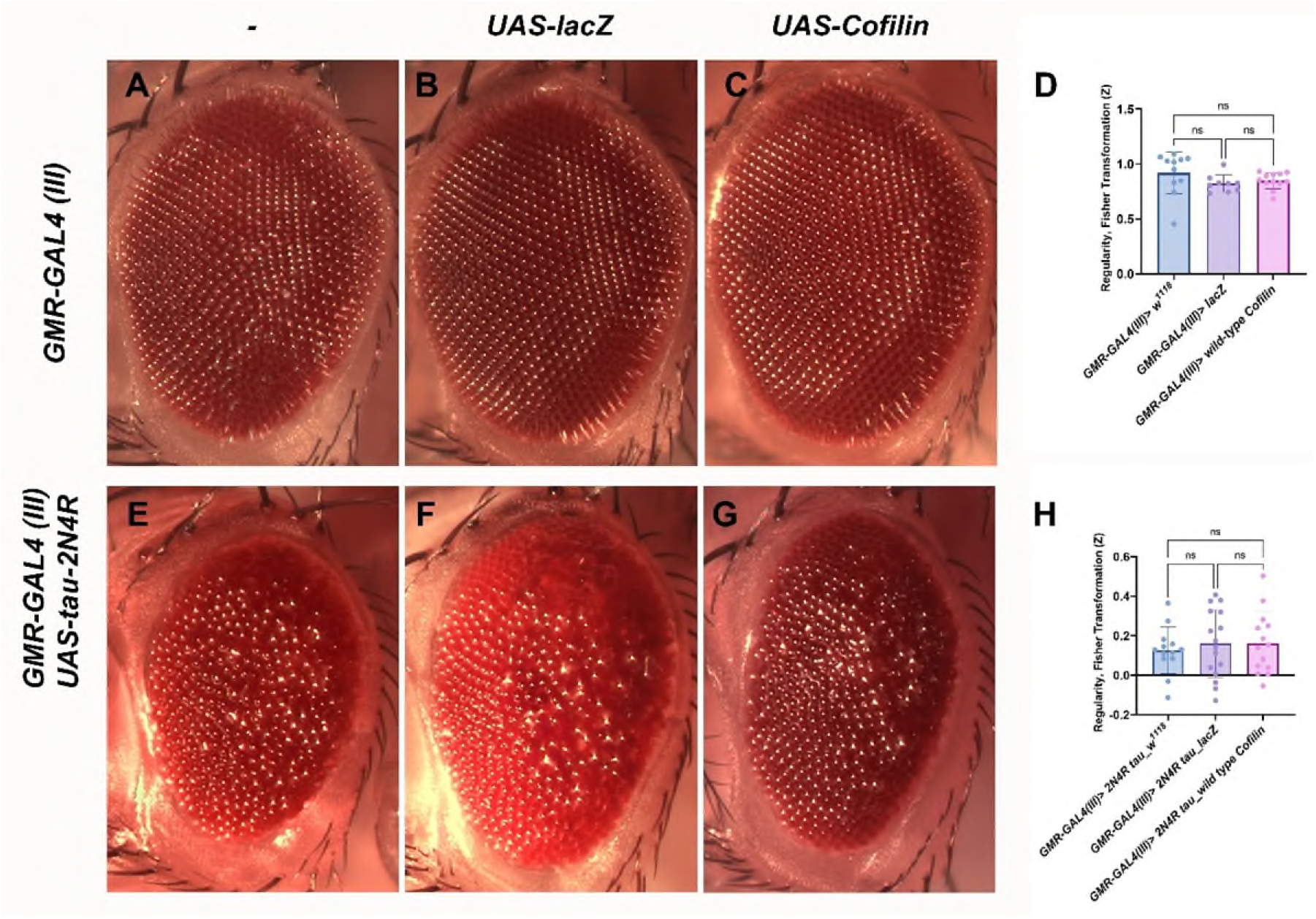
The rescue of htau-induced morphological eye defects by Cofilin overexpression is not detected using an automated assay for analysis of ommatidial regularity. **A-C.** Stereomicroscopic images of eyes from 2-4-day-old females carrying the third chromosome *GMR-GAL4 (III)* insertion in the presence of no other transgene (**A**), *UAS-lacZ* (**B**) or *UAS-Cofilin* (**C**). **D.** Bar chart plotting regularity score (as a Fisher Z transformation) for eyes of these genotypes relative to control *GMR-GAL4 (III)* reference curve reveals no detectable independent effect of expressing these transgenes. **E-G.** Stereomicroscopic images of eyes from 2-4-day-old females carrying the second chromosome *GMR-GAL4 (II)* insertion in the presence of *UAS-htau-2N4R* and co-expressing either no other transgene (**E**), *UAS-lacZ* (**F**) or *UAS-Cofilin* (**G**). **H.** Bar chart plotting regularity score (as a Fisher Z transformation) for eyes of these genotypes relative to control *GMR-GAL4 (II)* reference curve. The rescue of htau-induced morphological disruption in the eye by Cofilin overexpression (Fulga et al., 2007) is not detected using this assay, although the phenotype in the centre of the eye does appear to be partially suppressed. Scale bar: 100 µm.

**Fig. S3.**
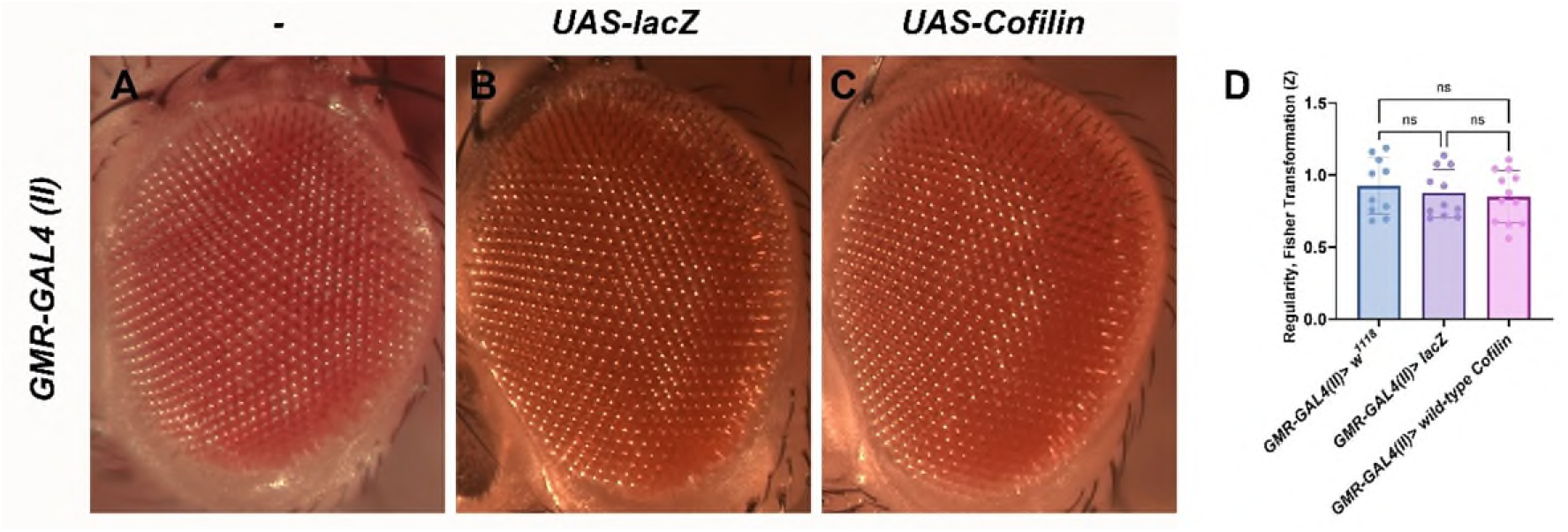
Expression of Cofilin with the GMR-GAL4 (II) driver does not induce morphological disruption in the eye. **A-C.** Stereomicroscopic images of eyes from 2-4-day-old females carrying the second chromosome *GMR-GAL4 (II)* insertion in the presence of no other transgene (**A**), *UAS-lacZ* (**B**) or *UAS-Cofilin* (**C**). **D.** Bar chart plotting regularity score (as a Fisher Z transformation) for eyes of these genotypes relative to control *GMR-GAL4 (II)* reference curve reveals no detectable independent effect of expressing these transgenes.

**Table S1.**
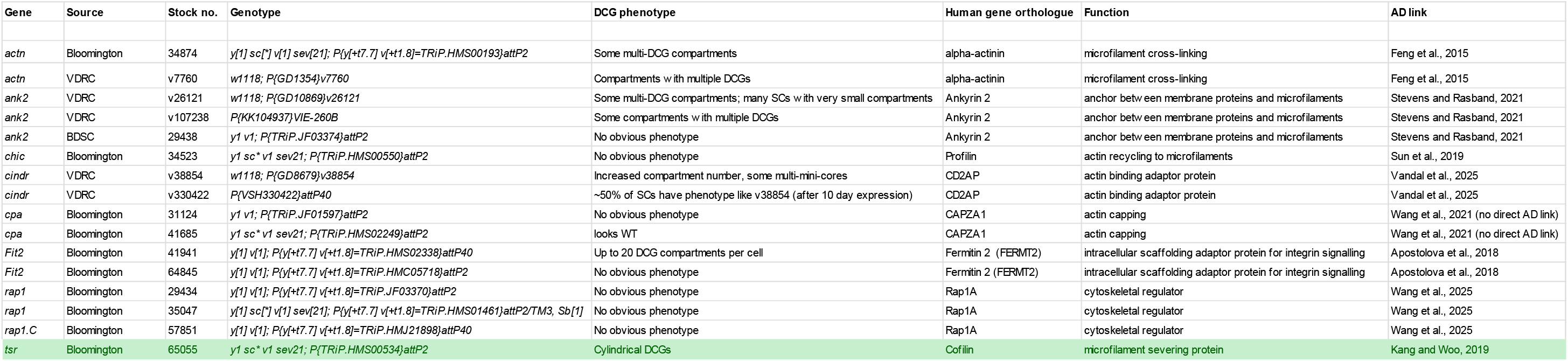
RNAi screen of microfilament cytoskeletal regulators for effects on DCG biogenesis. Findings from small-scale RNAi screen of microfilament cytoskeletal regulators, including genes encoding junctional proteins, which have been implicated in AD. Table details UAS-RNAi lines employed, human gene orthologue, cytoskeletal functions and references linking genes to AD, as well as any potential DCG phenotype. Although several genetic manipulations altered the number of DCG compartments or DCG structure, suggesting an important role for the actin cytoskeleton in DCG compartment maturation, only knockdown of *tsr* (*Drosophila cofilin*; shaded green) induced a cylindrical DCG phenotype that mirrored the effects of htau overexpression.

